# CAD rewires nucleotide metabolism to drive oxidative stress in myocardial ischemia/reperfusion injury

**DOI:** 10.64898/2026.05.15.725578

**Authors:** Xiaoshuai Zhao, Shuting Xu, Fangchi Yu, Yan Shi, Mengying Jin, Yuxuan Zhang, Kaichao Zhang, Jian Wang, Fan Zhang, Ying Liu, Junfang Wu, Guangyu Zhang, Xiaoding Wang

## Abstract

Myocardial ischemia/reperfusion (I/R) injury is a major determinant of infarct size and clinical outcome, yet effective therapies remain limited. Although metabolic remodeling is central to I/R pathology, the contribution of nucleotide biosynthesis remains unclear. Here, we identify CAD, the multifunctional rate-limiting enzyme of de novo pyrimidine biosynthesis, as a previously unrecognized regulator of myocardial reperfusion injury. CAD activation exacerbated cardiomyocyte death during simulated I/R, whereas CAD knockdown or pharmacological inhibition was protective in vitro. Mechanistically, CAD enhanced dihydroorotate dehydrogenase (DHODH)-dependent electron transfer, increased the CoQH_2_/CoQ ratio, and promoted complex I reverse electron transport (RET), thereby amplifying mitochondrial ROS. In parallel, CAD suppressed de novo purine synthesis, causing purine insufficiency, DIS3L-dependent RNA decay, and cytosolic ROS. Importantly, cardiomyocyte-specific CAD deletion protected against cardiac I/R injury in vivo. Together, these findings establish CAD as a metabolic hub linking nucleotide flux to dual-compartment ROS signaling and identify nucleotide metabolism as a therapeutic vulnerability in myocardial I/R injury.

**Graphical Abstract:** 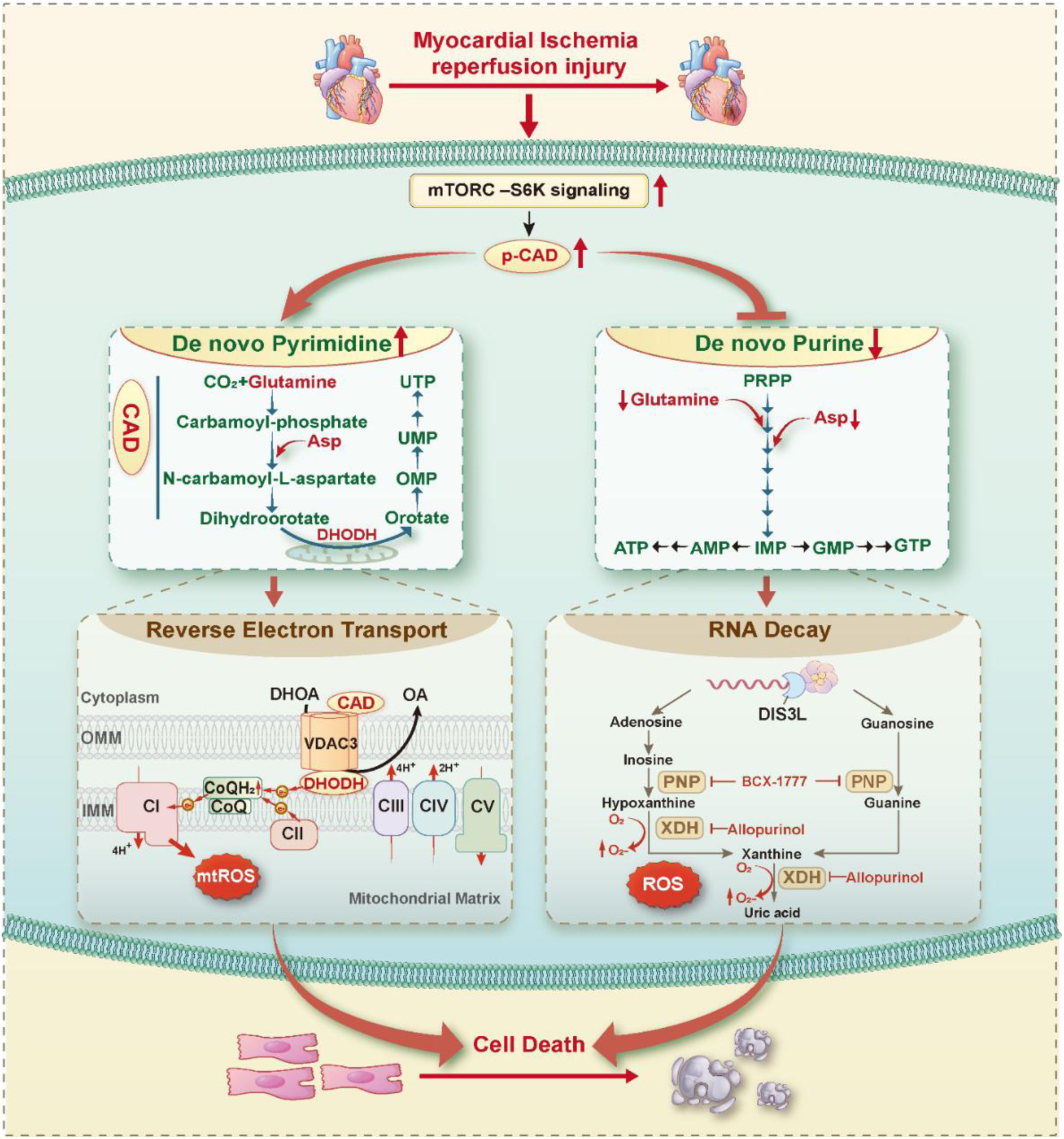

## Introduction

Ischemic heart disease remains the leading cause of morbidity and mortality worldwide, and timely restoration of blood flow is the cornerstone of therapy for acute myocardial infarction.^1–5^ However, although reperfusion salvages viable myocardium, the abrupt reintroduction of oxygen and nutrients paradoxically causes additional injury. This reperfusion phase is characterized by bursts of reactive oxygen species (ROS), calcium overload, mitochondrial permeability transition, and activation of regulated cell death programs, all of which substantially influence infarct size and post-ischemic remodeling.^6–10^ Despite its major clinical importance, effective therapies that directly target myocardial ischemia/reperfusion (I/R) injury remain limited.

A defining feature of myocardial ischemia and early reperfusion is profound metabolic remodeling. During ischemia, restricted oxygen availability suppresses mitochondrial oxidative metabolism and shifts ATP production toward anaerobic glycolysis. Upon reperfusion, rapid restoration of oxygen and substrates reactivates mitochondrial respiration and promotes ROS generation.^11–13^ In particular, reverse electron transport (RET) at mitochondrial complex I has emerged as a major source of reperfusion-associated ROS. In this model, ischemic succinate accumulation and a highly reduced coenzyme Q (CoQ) pool drive electron flow backward through complex I during reperfusion, generating superoxide that promotes mitochondrial dysfunction and cell death.^7,14^ While this framework has focused primarily on TCA cycle intermediates and electron transport chain dynamics, how broader biosynthetic pathways contribute to ROS generation during I/R remains incompletely understood.

*De novo* pyrimidine biosynthesis is initiated by the multifunctional enzyme CAD (carbamoyl-phosphate synthetase 2, aspartate transcarbamoylase, and dihydroorotase), followed by DHODH and cytosolic uridine monophosphate synthase (UMPS). Genetic defects in pyrimidine biosynthetic enzymes cause severe developmental and metabolic disorders, and pharmacologic targeting of this pathway has shown therapeutic value in cancer and immune diseases with high biosynthetic demand.^15–19^ Here, we identify CAD activation as a previously unrecognized metabolic driver of myocardial reperfusion injury. We demonstrate that CAD as a metabolic hub that couples nucleotide metabolism rewiring to compartment-specific ROS generation and cell death in myocardial reperfusion injury. By uncovering a previously unappreciated role for *de novo* pyrimidine metabolism in cardiac I/R, this work expands current paradigms of ischemic reperfusion injury and identifies CAD and its downstream nucleotide stress axis as promising therapeutic targets for ischemic heart disease.

## Results

### Cardiac ischemia/reperfusion induces de novo pyrimidine biosynthesis and activates CAD via mTOR–S6K signaling

Metabolic remodeling is a defining feature of cardiac ischemia/reperfusion (I/R), but the specific pathways involved during early reperfusion remain incompletely characterized. To systematically characterize reperfusion-associated metabolic alterations, we performed untargeted metabolomic profiling of ischemic myocardial regions from wild-type (WT) mice subjected to 45 min ischemia followed by 0 min, 30 min, or 4 h reperfusion, compared with sham controls. Heatmap analysis revealed selective accumulation of metabolites in the *de novo* pyrimidine biosynthetic pathway, including N-carbamoyl-L-aspartate and orotate, particularly at 30 min reperfusion (Figure 1A). Consistent with this observation, metabolite set enrichment analysis identified pyrimidine metabolism as the most prominently remodeled pathways at this time point (Figure 1B), indicating activation of *de novo* pyrimidine synthesis during early reperfusion.

**Figure 1.**
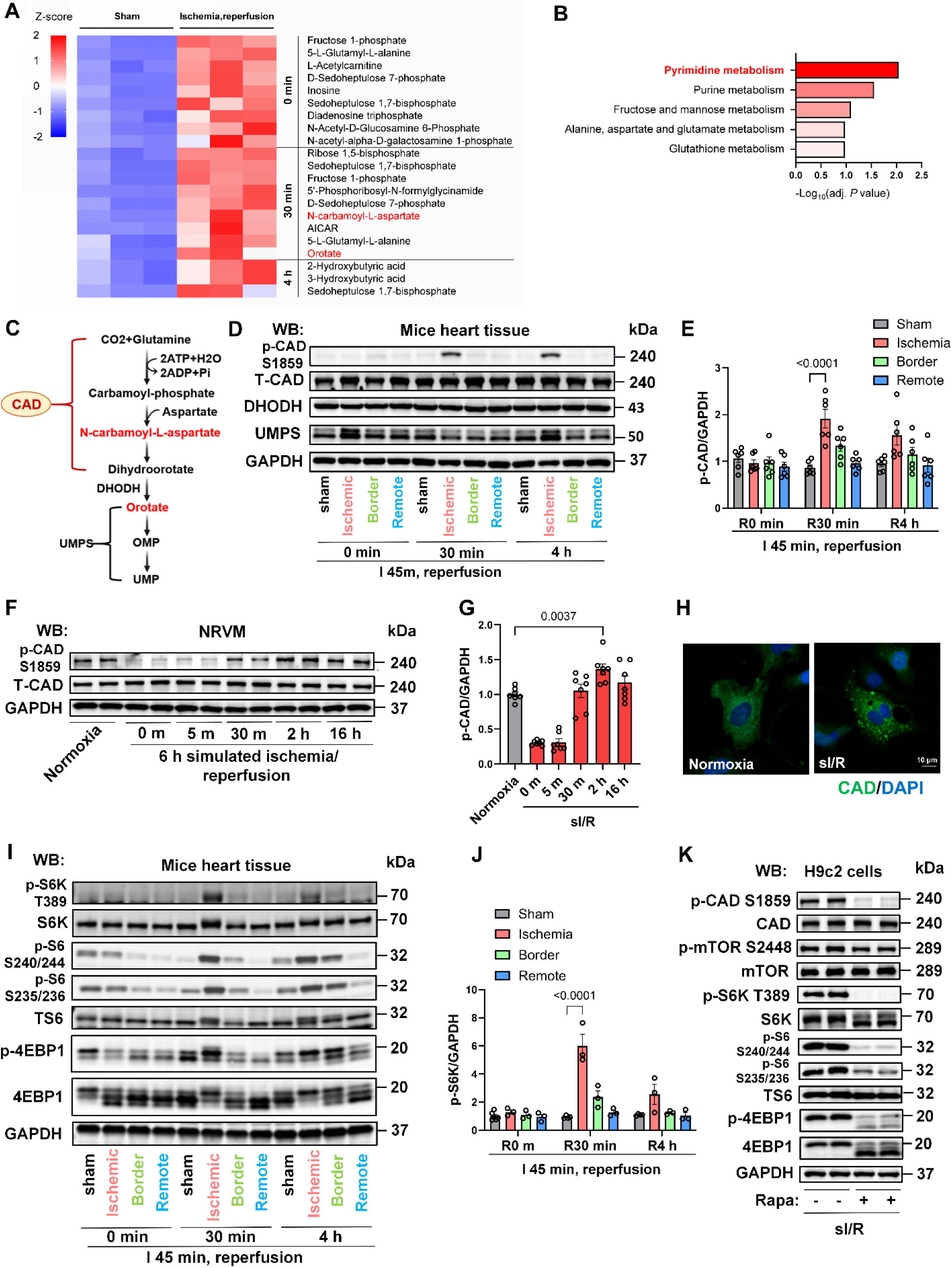
Cardiac ischemia/reperfusion activates CAD and de novo pyrimidine metabolism (A) Heatmap showing relative metabolite abundance in ischemic regions of wild-type (WT) mouse hearts under sham conditions or after 45 min cardiac ischemia followed by 0 min, 30 min, or 4 h reperfusion. n = 3. (B) Metabolite set enrichment analysis comparing WT sham hearts and WT hearts after ischemia/reperfusion (I/R) with 30 min reperfusion. (C) Schematic of the de novo pyrimidine biosynthetic pathway. (D) Immunoblot analysis of phosphorylated CAD (p-CAD), total CAD, DHODH, and UMPS in mouse hearts collected at the indicated reperfusion times (0, 30 min, and 4 h). Hearts were separated into ischemic, border, and remote regions. (E) Quantification of (D). n = 6. (F) Immunoblot analysis of p-CAD and total CAD in NRVMs subjected to normoxia or sI/R for the indicated reperfusion durations (0, 5 min, 30 min, 2 h, and 16 h). (G) Quantification of (F). n = 7. (H) Immunocytochemical staining of endogenous CAD in NRVMs under normoxic conditions or following sI/R for 2 h. Scale bars, 10 μm. (I) Immunoblot analysis of mTOR signaling pathway components in mouse hearts at indicated reperfusion time points (0, 30 min, and 4 h). (J) Quantification of (I). n = 3–6. (K) Immunoblot analysis of p-CAD, total CAD, and mTOR signaling components in H9c2 cells subjected to sI/R for 2 h in the presence of vehicle or rapamycin (10 nM). Data are presented as mean ± SEM. Statistical analyses were performed using two-way ANOVA followed by Tukey’s post hoc tests (E, J) or one-way ANOVA followed by Dunnett’s multiple-comparisons test (G). WB, Western blot.

We next examined whether the enzymatic machinery controlling *de novo* pyrimidine biosynthesis is dynamically regulated during I/R (Figure 1C). Immunoblot analysis revealed a robust increase in phosphorylation of CAD, the multifunctional enzyme catalyzing the first three committed steps of pyrimidine synthesis, in ischemic myocardial regions during reperfusion, whereas total CAD, DHODH, and UMPS protein levels remained unchanged across time points (Figures 1D and 1E). Similar regulation was observed *in vitro*: neonatal rat ventricular myocytes (NRVMs) and H9c2 cells subjected to simulated ischemia/reperfusion (sI/R) displayed significantly increased CAD phosphorylation during reperfusion without changes in total CAD abundance (Figures 1F and 1G; Figures S1A and S1B). In parallel, immunofluorescence analysis revealed a striking redistribution of endogenous CAD from a diffuse cytosolic pattern under normoxia to discrete punctate structures following sI/R (Figure 1H; Figure S1C). This reorganization is consistent with activity-dependent CAD assembly and supports functional activation of the *de novo* pyrimidine pathway.

Given prior evidence that CAD phosphorylation at Ser1859 is mediated by mTORC1–S6K signaling,^20,21^ we next examined activation of this upstream pathway. Components of the mTOR signaling cascade were activated in I/R myocardium and in sI/R-treated cardiomyocytes (Figures 1I and 1J; Figures S1D and S1E). Importantly, pharmacologic inhibition of mTORC1 with rapamycin or S6K with PF-4708671 markedly suppressed sI/R-induced CAD phosphorylation (Figure 1K; Figure S1F), demonstrating that CAD activation during reperfusion is mediated by mTOR–S6K signaling.

Together, these findings identify early reperfusion as a state of active *de novo* pyrimidine metabolic engagement in the heart. CAD is rapidly activated through mTORC1–S6K-dependent phosphorylation and undergoes dynamic subcellular reorganization, establishing CAD as a previously underappreciated metabolic node engaged during myocardial I/R injury.

### Cardiomyocyte CAD is necessary and sufficient to modulate ischemia/reperfusion injury *in vitro*

We next asked whether CAD regulates susceptibility to reperfusion injury in a cardiomyocyte-autonomous manner. We silenced CAD in H9c2 cells and neonatal rat ventricular myocytes (NRVMs) using siRNA (Figure 2A). CAD knockdown significantly reduced cell death induced by sI/R and H_2_O_2_ treatment in both H9c2 cells and NRVMs (Figures 2B and 2C). SYTOX Green staining further confirmed the protective effect of CAD silencing (Figure 2D). Pharmacological inhibition of CAD with N-(phosphonacetyl)-L-aspartate (PALA) similarly attenuated sI/R-induced cell death (Figure 2E), indicating that CAD enzymatic activity contributes to reperfusion-associated injury. Together, these data demonstrate that CAD loss or inhibition protects cardiomyocytes from oxidative and reperfusion stress *in vitro*.

**Figure 2.**
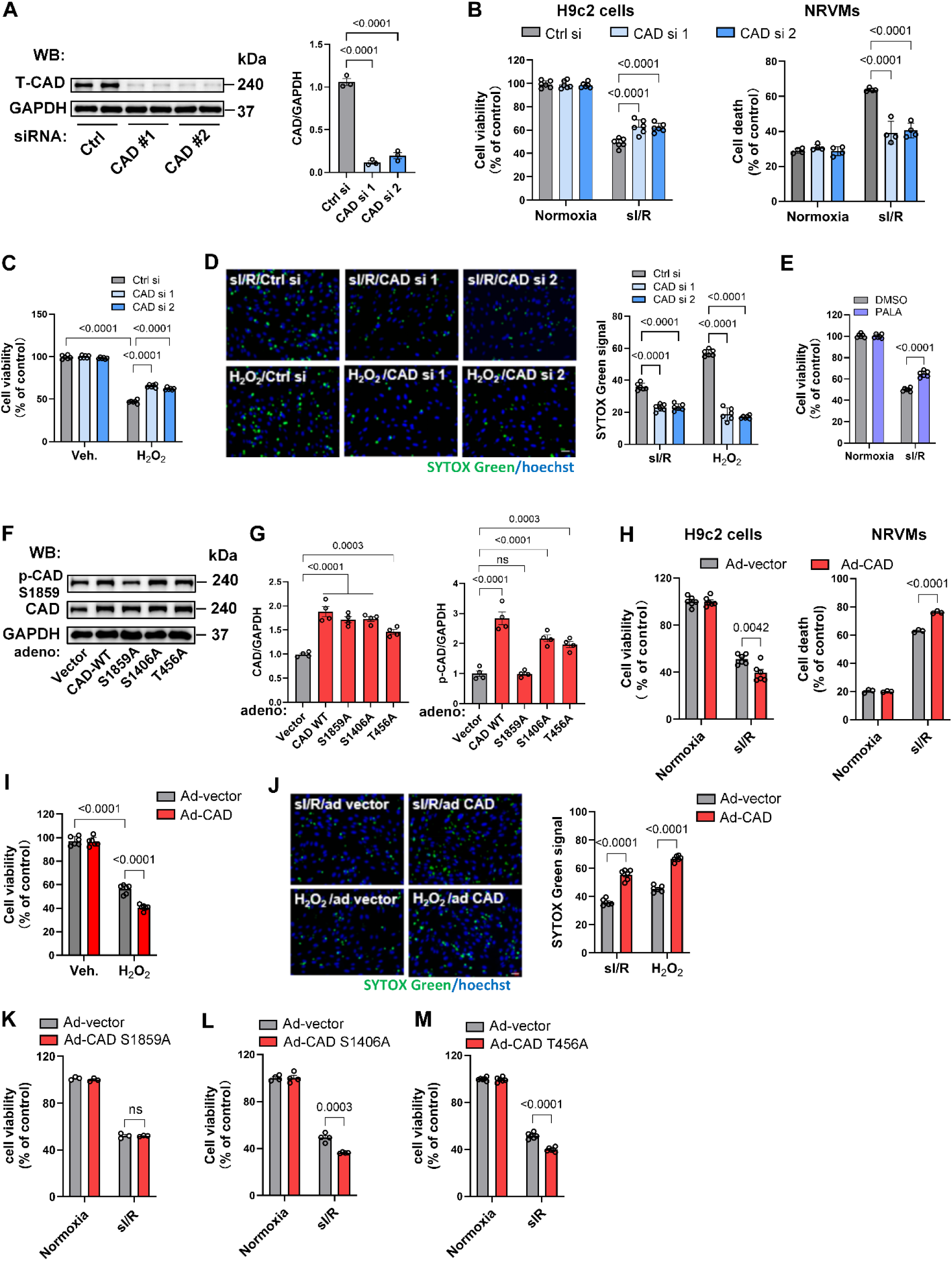
Cardiomyocyte CAD loss and activation bidirectionally regulate ischemia/reperfusion injury *in vitro* (A) Immunoblot validation of siRNA-mediated CAD knockdown in H9c2 cells and quantification of CAD knockdown efficiency. n = 3. (B) CAD knockdown significantly improves cell survival following sI/R in H9c2 cells (n = 6) and reduces cell death in NRVMs (n = 4). (C) CAD knockdown significantly improves cell survival in H9c2 cells subjected to H_2_O_2_–induced oxidative stress. n = 6. (D) Representative SYTOX Green and Hoechst staining images showing reduced cell death in H9c2 cells transfected with two independent CAD siRNAs compared with control siRNA following sI/R or H_2_O_2_ exposure. Quantification of SYTOX Green-positive cells normalized to total Hoechst-positive nuclei is shown. Scale bar, 200 μm. n = 6. (E) Pharmacological inhibition of CAD with PALA (10 μM) protects H9c2 cells from sI/R-induced injury. n = 6. (F) Immunoblot analysis of CAD overexpression and phosphorylation-defective mutants. Adenoviral expression of CAD-WT and indicated mutants increases total CAD protein abundance. CAD-WT, CAD-S1406A, and CAD-T456A increase Ser1859 phosphorylation, whereas CAD-S1859A does not. (G) Quantification of (F). n = 4. (H) CAD overexpression exacerbates sI/R-induced injury in H9c2 cells (n = 6) and NRVMs (n = 3). (I) CAD overexpression exacerbates H_2_O_2_-induced injury, reducing cell viability in H9c2 cells. n = 6. (J) Representative SYTOX Green and Hoechst staining images showing increased cell death in CAD-overexpressing cells compared with vector controls following sI/R or H_2_O_2_ exposure. Quantification of SYTOX Green-positive cells normalized to total Hoechst-positive nuclei is shown. Scale bar, 200 μm. n = 6. (K-M) Functional analysis of CAD phosphorylation mutants. Unlike CAD-WT, CAD-S1859A fails to exacerbate sI/R-induced injury, whereas CAD-S1406A and CAD-T456A retain this effect. n = 3–6. Data are presented as mean ± SEM. Statistical analyses were performed using one-way ANOVA followed by Dunnett’s multiple-comparisons test (A, G) or two-way ANOVA followed by Tukey’s or Sidak’s post hoc tests (B–E, H–M). WB, western blot; ns, not significant.

We next used gain-of-function approaches to determine whether CAD activation is sufficient to exacerbate injury. Because CAD activity is regulated by phosphorylation at multiple residues, including Ser1859, Ser1406, and Thr456, we generated adenoviruses expressing CAD-WT or phosphorylation-defective mutants. Expression of CAD-WT, CAD-S1406A, and CAD-T456A markedly increased CAD protein abundance and retained Ser1859 phosphorylation, whereas the CAD-S1859A mutant failed to increase Ser1859 phosphorylation (Figures 2F and 2G). CAD overexpression significantly exacerbated cell death in response to sI/R and H_2_O_2_ treatment (Figures 2H and 2I). Consistent with these findings, SYTOX Green staining demonstrated increased fluorescence intensity in CAD-overexpressing cells, indicating enhanced cell death (Figure 2J). Importantly, the S1859A mutant failed to promote sI/R-induced cell death, whereas the S1406A and T456A mutants remained fully competent to exacerbate injury (Figures 2K–2M). These findings demonstrate that Ser1859 phosphorylation is specifically required for CAD-mediated sensitization to ischemia/reperfusion injury.

### CAD induction rewires nucleotide metabolism during reperfusion

Having established that CAD activation promotes reperfusion injury, we next sought to define the underlying metabolic alterations. Given that CAD catalyzes the first committed steps of de novo pyrimidine biosynthesis and thereby occupies a central position in nucleotide metabolism, we reasoned that its activation may directly perturb nucleotide metabolism during reperfusion. Targeted metabolomic analysis in CAD-overexpressing H9c2 cells subjected to sI/R revealed robust accumulation of N-carbamoyl-L-aspartate, dihydroorotate (DHOA), and orotate, together with a modest increase in UMP, in CAD-overexpressing cells under sI/R conditions (Figures 3A–3D). In contrast, purine nucleotides, including IMP, AMP, and GMP, were significantly reduced (Figures 3E–3G), indicating a shift in nucleotide balance toward pyrimidine synthesis.

**Figure 3.**
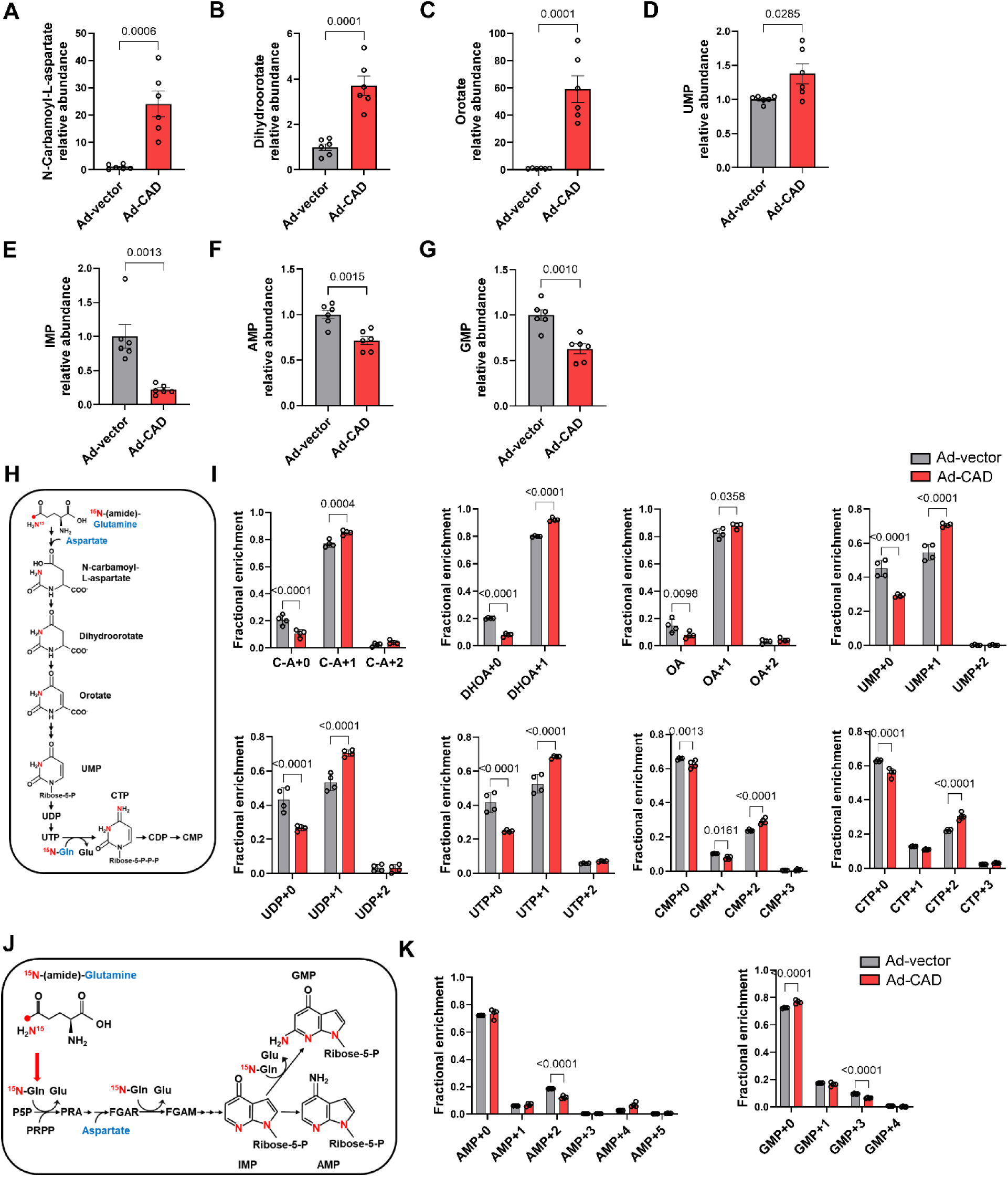
CAD reprograms nucleotide metabolism by enhancing de novo pyrimidine synthesis and suppressing purine biosynthesis. (A–D) Targeted metabolomic analysis showing increased levels of de novo pyrimidine intermediates, including N-carbamoyl-L-aspartate (C-A) (A), dihydroorotate (DHOA) (B), orotate (OA) (C), and UMP (D), in CAD-overexpressing H9c2 cells subjected to sI/R. n = 6. (E–G) Targeted metabolomic analysis showing decreased levels of purine nucleotides, including IMP (E), AMP (F), and GMP (G), in CAD-overexpressing H9c2 cells following sI/R. n = 6. (H) Schematic of [amide-^15^N]glutamine tracing strategy used to assess de novo pyrimidine biosynthesis. (I) Fractional enrichment analysis demonstrating increased ^15^N incorporation into pyrimidine intermediates and nucleotides in CAD-overexpressing H9c2 cells following sI/R. n = 4. (J) Schematic of the [amide-^15^N]glutamine tracing strategy used to assess de novo purine biosynthesis. (K) Fractional enrichment analysis showing reduced ^15^N incorporation into purine nucleotides in CAD-overexpressing cells following sI/R. n = 4. Data are presented as mean ± SEM. Statistical analyses were performed using two-sided unpaired Student’s t-tests (A-G) or two-way ANOVA followed by Tukey’s or Sidak’s post hoc tests (I, K).

Because *de novo* pyrimidine and *de novo* purine biosynthetic pathways share key substrates, including glutamine and aspartate, we hypothesized that CAD activation redirects metabolic flux toward pyrimidine synthesis at the expense of purine production. To test this, we performed stable isotope tracing using [amide-^15^N]glutamine in vector control and CAD-overexpressing H9c2 cells subjected to sI/R. CAD overexpression markedly increased ^15^N in comparison to incorporation into pyrimidine intermediates and nucleotides, consistent with enhanced flux through the *de novo* pyrimidine pathway (Figures 3H and 3I; Figure S2A). In contrast, ^15^N incorporation into purine nucleotides was reduced, indicating suppression of *de novo* purine biosynthetic flux (Figures 3J and 3K; Figures S2B-S2E). Together, these data demonstrate that CAD activation during reperfusion reprograms nucleotide metabolism by driving *de novo* pyrimidine synthesis while constraining purine biosynthesis.

### CAD activation drives DHODH-dependent reverse electron transport and mitochondrial ROS production

Because CAD activation markedly increased DHOA and OA, we next asked whether enhanced flux through the CAD–DHODH arm of *de novo* pyrimidine biosynthesis alters mitochondrial redox state. Immunoblot analysis showed that DHODH and UMPS protein abundance remained unchanged in CAD-overexpressing cells relative to controls under normoxia or sI/R conditions (Figure S3A), indicating that the downstream effects reflect altered pathway flux rather than increased enzyme abundance. DHODH oxidizes DHOA to OA while transferring electrons to the CoQ pool (Figure 4A).^22–24^ Consistent with enhanced electron transfer through DHODH, CAD overexpression increased the CoQH_2_/CoQ ratio following sI/R, particularly at later reperfusion time points (Figure 4B), indicating a more reduced CoQ pool that is permissive for reverse electron transport.

**Figure 4.**
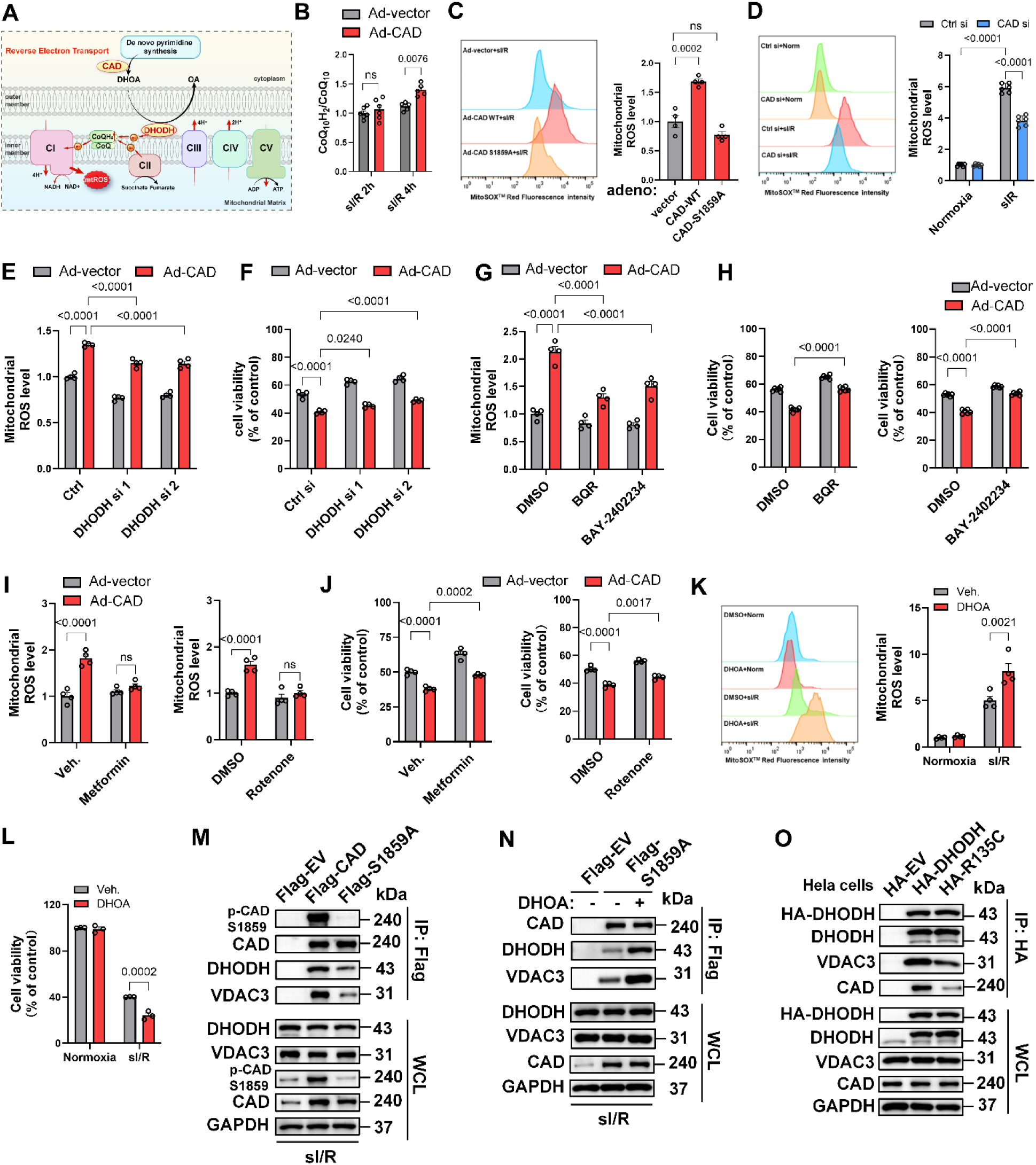
CAD activation drives DHODH-dependent reverse electron transport and mitochondrial ROS production. (A) Schematic illustrating how DHODH transfers electrons to the CoQ pool, increasing CoQH_2_ levels and promoting complex I reverse electron transport (RET)–dependent mitochondrial ROS generation during reperfusion. (B) Quantification of the CoQH_2_/CoQ ratio in CAD-overexpressing H9c2 cells following sI/R for 2 h or 4 h. n = 5–6. (C) Representative overlay histograms and quantification of MitoSOX fluorescence in H9c2 cells following sI/R for 4 h. CAD overexpression shifts fluorescence intensity rightward, whereas CAD-S1859A does not. n = 4. (D) Representative overlay histograms and quantification of MitoSOX fluorescence in H9c2 cells under normoxia or following sI/R for 4 h. CAD knockdown shifts fluorescence intensity leftward and reduces mitochondrial ROS. n = 6. (E) DHODH silencing partially rescues CAD-induced mitochondrial ROS elevation after sI/R. n = 4. (F) DHODH knockdown partially rescues CAD-induced cell death during sI/R. n = 4. (G) Pharmacologic inhibition of DHODH with brequinar (BQR, 10 nM) or BAY-2402234 (10 nM) reduces mitochondrial ROS production in CAD-overexpressing H9c2 cells following sI/R. n = 4. (H) DHODH inhibition with brequinar or BAY-2402234 partially restores cell viability in CAD-overexpressing cells subjected to sI/R. n = 6. |(I) Metformin (2 μM) or rotenone (5 nM) rescues mitochondrial ROS production in CAD-overexpressing H9c2 cells following sI/R. n = 4. (J) Metformin (2 μM) or rotenone (5 nM) partially rescues cell viability in CAD-overexpressing H9c2 cells subjected to sI/R. n = 4. (K) Representative overlay histograms and quantification of MitoSOX fluorescence in H9c2 cells under normoxia or following sI/R. DHOA (50 μM) further increases mitochondrial ROS during sI/R and shifts fluorescence intensity rightward. n = 4. (L) DHOA (50 μM) increases sI/R-induced cell death in H9c2 cells. n = 3. (M) Co-immunoprecipitation in H9c2 cells expressing Flag-EV, Flag-CAD, or Flag-CAD-S1859A showing interaction of CAD with DHODH and VDAC3. (N) Co-immunoprecipitation in H9c2 cells expressing Flag-CAD-S1859A with or without DHOA showing restoration of CAD-DHODH-VDAC3 complex formation by DHOA. (O) Co-immunoprecipitation in HeLa cells expressing HA-EV, HA-DHODH, or HA-DHODH-R135C showing interaction of DHODH with CAD and VDAC3. Data are presented as mean ± SEM. Statistical analyses were performed using two-way ANOVA followed by Tukey’s post hoc tests (B, D-L) or one-way ANOVA followed by Dunnett’s multiple-comparisons test (C). WB, Western blot. ns denotes not significant.

Accordingly, CAD overexpression markedly increased mitochondrial ROS following sI/R, whereas the phosphorylation-defective S1859A mutant failed to do so (Figure 4C). Conversely, CAD knockdown significantly reduced mitochondrial ROS accumulation (Figure 4D), and pharmacological inhibition of CAD with PALA similarly suppressed mitochondrial ROS production (Figure S3B), indicating that CAD enzymatic activity is required for this process. To directly assess the role of DHODH, we silenced DHODH expression in H9c2 cells (Figures S3C and S3D). DHODH knockdown partially rescued CAD-induced mitochondrial ROS elevation and cell viability during sI/R (Figures 4E and 4F; Figure S3E). Consistently, pharmacological inhibition of DHODH with brequinar (BQR) or BAY-2402234 reduced mitochondrial ROS and improved cell survival in CAD-overexpressing cells (Figures 4G and 4H; Figure S3F), demonstrating that DHODH activity is required for CAD-driven mitochondrial oxidative injury.

To determine whether CAD-induced mitochondrial ROS arises from RET rather than forward electron leak, we treated cells with the complex I inhibitors metformin and rotenone,^25–28^ both of which suppress RET-derived ROS. These interventions significantly reduced mitochondrial ROS and improved cell viability in CAD-overexpressing cells subjected to sI/R (Figures 4I and 4J; Figures S3G and 3H), supporting a RET-dependent mechanism. Moreover, supplementation with DHOA further increased mitochondrial ROS and exacerbated cell death (Figures 4K and 4L), indicating that enhanced substrate supply to the CAD–DHODH axis is sufficient to amplify mitochondrial oxidative stress.

Previous studies have suggested that CAD associates with DHODH at the mitochondrial outer membrane through VDAC3.^23^ Consistent with this, co-immunoprecipitation demonstrated robust interactions among CAD, DHODH, and VDAC3, whereas the phosphorylation-defective CAD S1859A mutant exhibited markedly reduced association with DHODH and VDAC3 (Figure 4M; Figure S3I), indicating that CAD activation promotes assembly of this mitochondrial metabolic complex. Notably, supplementation with DHOA restored CAD–DHODH–VDAC3 complex formation in the S1859A context (Figure 4N), suggesting that substrate availability can stabilize this signaling complex. Consistently, wild-type DHODH, but not the catalytic mutant R135C, efficiently associated with CAD and VDAC3 in HeLa cells (Figure 4O).

Together, these data indicate that CAD activation enhances DHOA production, stimulates DHODH-dependent electron transfer to CoQ, promotes complex I reverse electron transport, and drives excessive mitochondrial ROS generation, culminating in cardiomyocyte death during reperfusion.

### CAD activation couples purine insufficiency to DIS3L-dependent RNA decay, cytosolic ROS, and cell death during reperfusion

We next investigated how CAD-mediated nucleotide rewiring contributes to cell injury during sI/R. Because balanced nucleotide pools are essential for RNA synthesis and stability,^29,30^ we hypothesized that CAD-driven suppression of purine availability may perturb RNA homeostasis.

To test this, we performed EU pulse–chase labeling to quantify global RNA synthesis and decay (Figure 5A). CAD overexpression did not affect nascent RNA synthesis, as reflected by comparable EU incorporation at the start of the chase (Figure 5B), but markedly accelerated RNA decay during reperfusion (Figure 5C). Conversely, CAD knockdown significantly slowed RNA decay without altering RNA synthesis (Figures 5D and 5E), demonstrating that CAD is both necessary and sufficient to promote RNA destabilization under sI/R.

**Figure 5.**
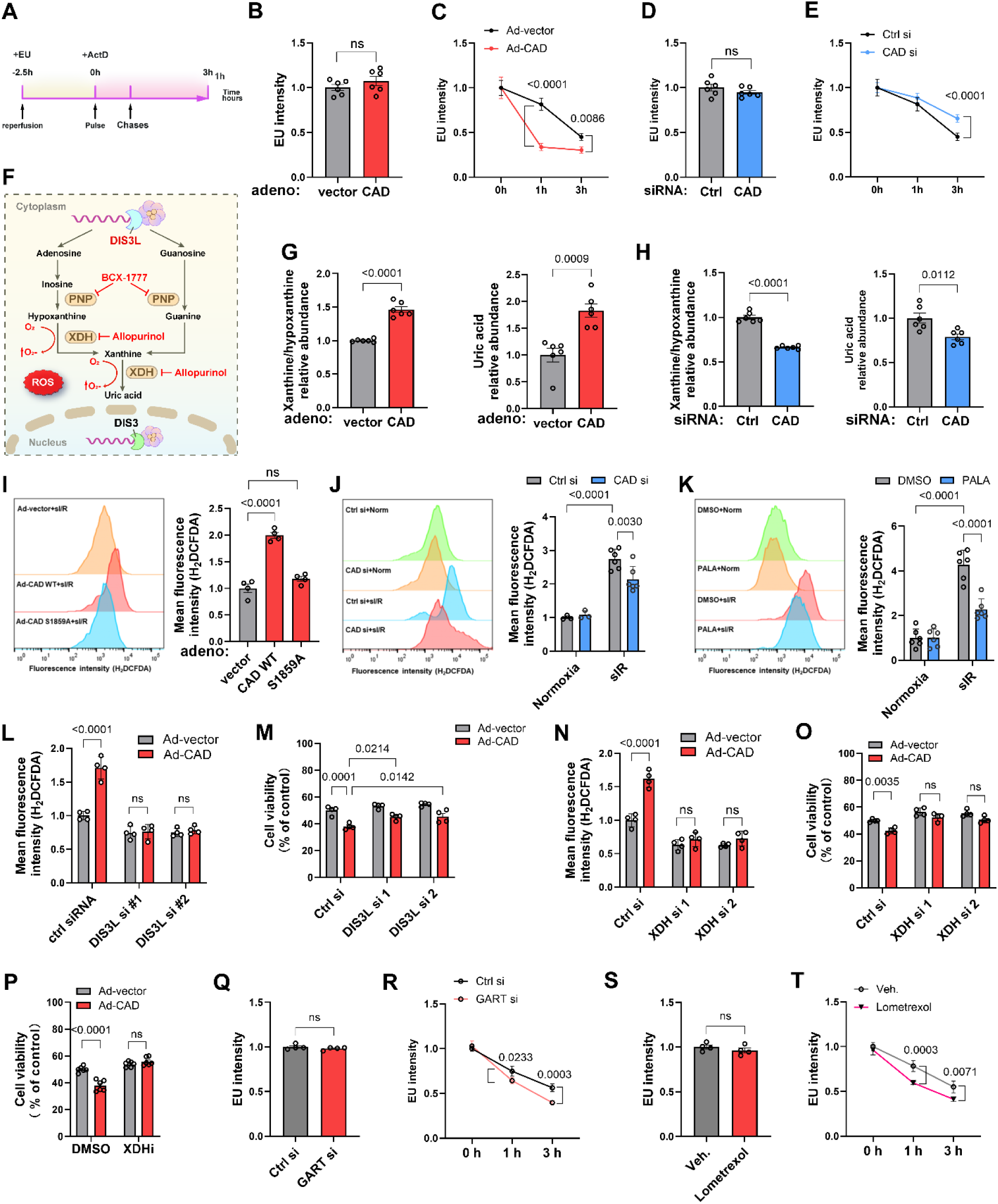
CAD activation drives DIS3L-dependent RNA decay, purine catabolism, and cytosolic ROS generation during reperfusion. (A) Schematic of the EU pulse-chase strategy used to assess RNA synthesis and decay. (B) Quantification of EU fluorescence intensity showing comparable RNA synthesis at 0 h. n = 6. (C) Quantification of accelerated RNA decay in CAD-overexpressing cells at later chase times. n = 6. (D) CAD knockdown does not alter EU incorporation. n = 6. (E) CAD knockdown significantly slows RNA decay following sI/R. n = 6. (F) Schematic of DIS3L-dependent cytoplasmic RNA decay and purine catabolism-linked ROS production. (G) CAD overexpression increases hypoxanthine/xanthine and uric acid accumulation during sI/R. n = 6. (H) CAD knockdown reduces hypoxanthine/xanthine and uric acid accumulation during sI/R. n = 6. (I) Representative overlay histograms and quantification of H_2_DCFDA fluorescence in H9c2 cells following sI/R. CAD overexpression shifts fluorescence intensity rightward, whereas CAD-S1859A does not. n = 4. (J) Representative overlay histograms and quantification of H_2_DCFDA fluorescence in H9c2 cells following sI/R. CAD knockdown shifts fluorescence intensity leftward and reduces cytosolic ROS. n = 6. (K) Representative overlay histograms and quantification of H_2_DCFDA fluorescence in H9c2 cells following sI/R. PALA (10 μM) shifts fluorescence intensity leftward and reduces cytosolic ROS. n = 6. (L) DIS3L silencing abolishes CAD-induced cytosolic ROS following sI/R. n = 4. (M) DIS3L knockdown partially rescues CAD-induced cell death following sI/R. n = 4. (N) XDH silencing abolishes CAD-induced cytosolic ROS during sI/R. n = 4. (O) XDH silencing rescues CAD-induced cell death following sI/R. n = 4. (P) Pharmacologic inhibition of XDH with allopurinol (20 μM) restores cell viability in CAD-overexpressing cells subjected to sI/R. n = 6. (Q and R) GART knockdown does not affect RNA synthesis at 0 h (Q) but accelerates RNA decay during the chase period (R). n = 4. (S and T) Lometrexol (5 μM) does not alter RNA synthesis at 0 h (S) but accelerates RNA decay following sI/R (T). n = 4. Data are presented as mean ± SEM. Statistical analyses were performed using two-sided unpaired Student’s *t*-tests (B, D, G, H, Q, S), two-way ANOVA followed by Tukey’s or Sidak’s post hoc tests (C, E, J, K, M, N, P, R, T), or one-way ANOVA followed by Dunnett’s multiple-comparisons test (I). WB, Western blot. ns denotes not significant.

Accelerated RNA degradation was accompanied by enhanced purine catabolism. CAD overexpression led to increased accumulation of hypoxanthine, xanthine, and uric acid during sI/R, whereas CAD knockdown reduced their levels (Figures 5G and 5H). Consistent with this, CAD activation increased cytosolic ROS in a phosphorylation-dependent manner, as the S1859A mutant failed to induce ROS, while CAD silencing or pharmacologic inhibition with PALA attenuated ROS accumulation (Figures 5I–5K).

To define the RNA decay machinery involved, we targeted components of the RNA exosome.^31–33^ Silencing DIS3L, the cytoplasmic 3’-5’ exoribonuclease responsible for bulk RNA degradation, attenuated RNA decay, abolished cytosolic ROS accumulation, and partially rescued cell viability in CAD-overexpressing cells (Figures 5L and 5M; Figure S4A–S4D). In contrast, knockdown of the nuclear exosome component DIS3 had no effect on RNA decay, ROS, or cell survival (Figures S4E–S4I), indicating that CAD-induced RNA degradation is mediated specifically through the cytoplasmic DIS3L pathway.

We next examined whether downstream purine catabolism is required for ROS production. Silencing xanthine dehydrogenase (XDH), pharmacologically inhibiting its activity with allopurinol, or blocking purine nucleoside phosphorylase (PNP) with BCX1777,^34^ abolished CAD-induced cytosolic ROS accumulation and CAD-mediated cell death (Figures 5N–5P; Figures S4J and S4L), establishing purine catabolism as the primary downstream source of ROS and injury following CAD activation..

Importantly, direct impairment of de novo purine synthesis was sufficient to phenocopy these effects. Genetic inhibition of GART or pharmacological inhibition with lometrexol did not affect RNA synthesis at baseline but markedly accelerated RNA decay during sI/R (Figures 5Q–5T; Figure S4M), demonstrating that purine insufficiency alone is sufficient to drive RNA destabilization independently of CAD overexpression.

Together, these findings define a pathway in which CAD activation suppresses purine availability, thereby triggering DIS3L-dependent RNA decay, fueling purine catabolism, and driving cytosolic ROS–mediated cardiomyocyte injury during reperfusion.

### Purine availability dictates RNA stability and oxidative stress during reperfusion

Given that CAD activation suppresses de novo purine biosynthesis, we next asked whether restoring purine supply could counteract RNA decay and downstream injury. Immunoblot analysis revealed upregulation of multiple de novo purine biosynthetic enzymes, including PPAT, GART, ADSL, and IMPDH1, in CAD-overexpressing cells during sI/R (Figures 6A and 6B; Figure S5A), suggesting a compensatory response to purine insufficiency. Notably, APRT1, a key enzyme in the purine salvage pathway^35^, was selectively induced under sI/R, whereas HPRT1 remained unchanged, indicating preferential activation of adenine salvage.

**Figure 6.**
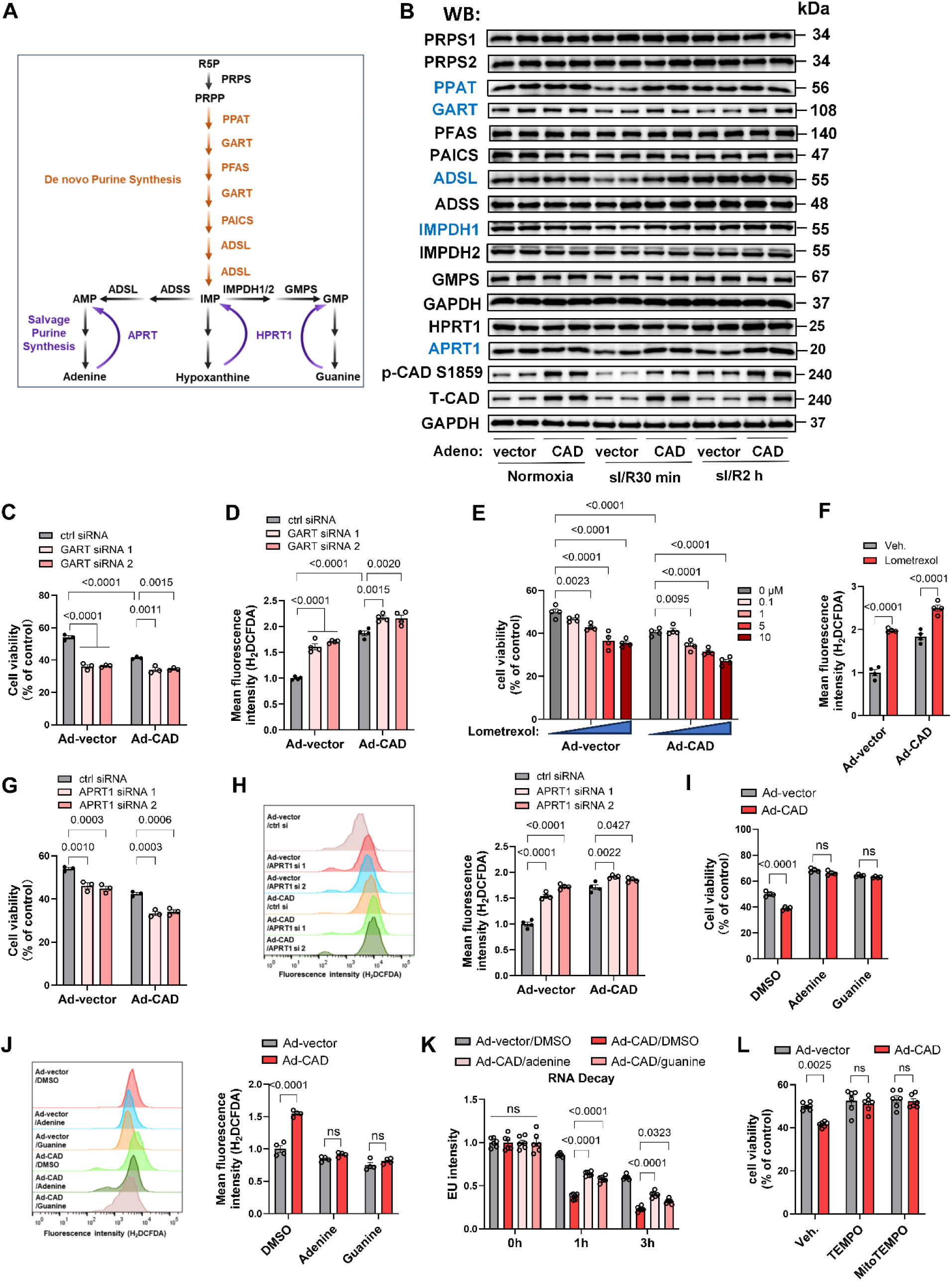
Purine depletion exacerbates, whereas salvage pathway restoration suppresses, CAD-driven RNA decay, ROS, and cell death. (A) Schematic illustrating de novo and salvage purine synthesis pathways. (B) Immunoblot analysis of purine biosynthetic enzymes, total CAD, and p-CAD in H9c2 cells expressing vector or CAD under normoxia or sI/R. (C) GART silencing exacerbates CAD-induced cell death following sI/R. n = 3. (D) GART silencing exacerbates CAD-induced cytosolic ROS accumulation. n = 4. (E) Dose-dependent exacerbation of CAD-induced cell death by pharmacological inhibition of GART with lometrexol (0–10 μM). n = 4. (F) Pharmacological inhibition of GART with lometrexol (5 μM) exacerbates CAD-induced cytosolic ROS accumulation. n = 3. (G and H) Silencing of APRT1 exacerbates CAD-induced cell death (G) and cytosolic ROS accumulation (H) in H9c2 cells subjected to sI/R. n = 3. (I-K) Supplementation with adenine (0.5 μM) or guanine (0.5 μM) rescues CAD-induced cell death (I), reduces cytosolic ROS accumulation (J), and attenuates accelerated RNA decay (K) following sI/R. n = 4–6. (L) Treatment with ROS scavengers (TEMPO and MitoTEMPO) restores cell viability in CAD-overexpressing cells following sI/R. n = 6. Data are presented as mean ± SEM. Statistical analyses were performed using two-way ANOVA followed by Tukey’s or Sidak’s post hoc tests (C-N). WB, Western blot. ns denotes not significant.

Functionally, disruption of purine synthesis exacerbated CAD-induced phenotypes. Silencing GART or pharmacologic inhibition with lometrexol increased cytosolic ROS accumulation and cell death under sI/R, with further exacerbation in CAD-overexpressing cells (Figures 6C–6F; Figures S5B-S5C). Similarly, silencing APRT1 aggravated CAD-induced cell death and ROS accumulation (Figures 6G and 6H; Figure S5D), demonstrating that both de novo and salvage pathways are required to maintain nucleotide homeostasis and limit injury.

Conversely, restoring purine availability robustly suppressed CAD-driven phenotypes. Supplementation with adenine or guanine rescued cell viability, reduced cytosolic ROS accumulation, and, notably, reversed the accelerated RNA decay induced by CAD activation (Figures 6I–6K). These findings establish purine availability as a key determinant of RNA stability under reperfusion stress.

Consistent with a ROS-dependent mechanism, treatment with ROS scavengers^36^ (TEMPO and MitoTEMPO) restored cell viability in CAD-overexpressing cells (Figure 6L). Together, these data support a model in which purine insufficiency drives RNA decay and oxidative stress, whereas restoration of purine pools suppresses these pathogenic processes.

### Cardiomyocyte-specific deletion of CAD protects against myocardial I/R injury *in vivo*

Having established that CAD drives cardiomyocyte injury through coordinated metabolic and redox mechanisms *in vitro*, we next asked whether loss of CAD protects the heart from ischemia/reperfusion injury *in vivo*. To this end, we generated a cardiomyocyte-specific CAD conditional knockout mouse model by crossing CAD^fl/fl^ mice with αMHC-Cre transgenic mice (CAD cKO; Figure 7A). Immunoblot analysis confirmed efficient deletion of CAD in cardiac tissue, while CAD expression in liver, kidney, and brain remained unchanged, demonstrating cardiac specificity of the knockout (Figure 7B; Figure S6A).

**Figure 7.**
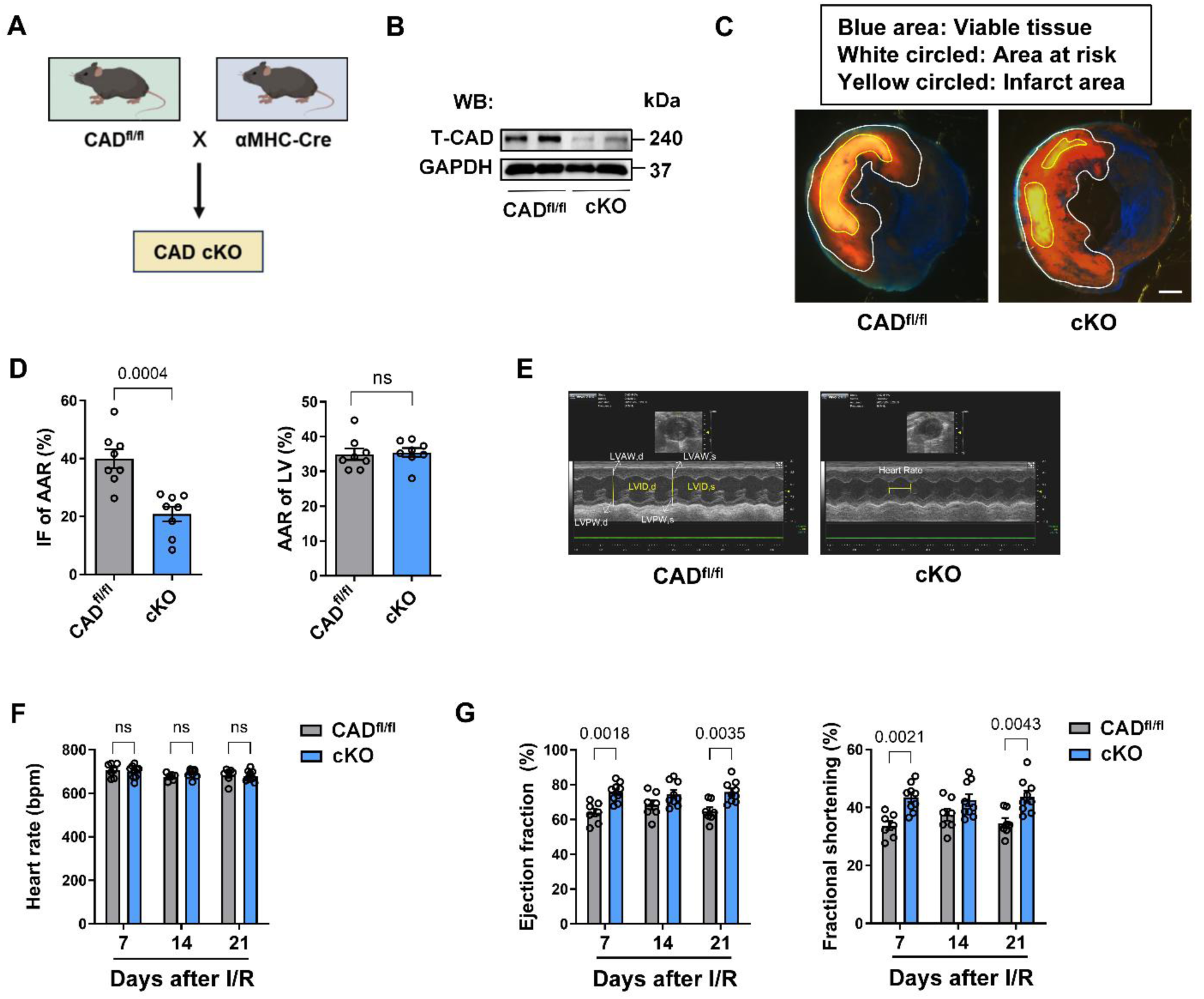
Cardiomyocyte CAD is required for myocardial ischemia/reperfusion injury. (A) Schematic of the cardiomyocyte-specific CAD conditional knockout mouse model. (B) Immunoblot validation of CAD deletion efficiency in CAD cKO mice. (C) CAD cKO mice and littermate controls were subjected to 45 min of cardiac ischemia followed by 24 h of reperfusion. Hearts were stained with TTC to delineate viable myocardium, area at risk (AAR), and infarcted tissue (IF). Representative images are shown; regions are outlined. Scale bar, 1 mm. (D) Quantification of infarct size expressed as IF/AAR and AAR/LV. CAD cKO mice exhibit significantly reduced IF/AAR, while AAR/LV is comparable between genotypes, indicating comparable I/R surgery. n = 8. (E) Representative echocardiographic images with labeled measurements. (F) Heart rate measurements following I/R in CAD^fl/fl^ and CAD cKO mice. Echocardiography was performed at 7, 14, and 21 days post-I/R. n = 7–9. (G) Echocardiographic assessment of cardiac function after I/R. Ejection fraction (EF) and fractional shortening (FS) were measured at 7, 14, and 21 days post-I/R, demonstrating improved functional recovery in CAD cKO mice compared with CAD^fl/fl^ controls. n = 7–9. Data are presented as mean ± SEM. Statistical analyses were performed using two-sided unpaired Student’s t-tests (D), two-way ANOVA followed by Tukey’s or Sidak’s post hoc tests (F, G). WB, Western blot. ns denotes not significant.

CAD cKO mice and littermate CAD^fl/fl^ controls were subjected to 45 min of coronary occlusion followed by 24 h reperfusion. Triphenyltetrazolium chloride (TTC) staining revealed a marked reduction in infarct size in CAD cKO hearts compared with CAD^fl/fl^ controls (Figures 7C and 7D). No difference in area at risk relative to left ventricle (AAR/LV) was observed, indicating comparable ischemic injury and surgical consistency between groups (Figure 7D). Consistent with reduced myocardial damage, echocardiographic assessment demonstrated significantly improved post-ischemic cardiac function in CAD cKO mice, with higher ejection fraction and fractional shortening at 7 and 21 days after I/R (Figures 7E–7G; Figures S6B and S6C).

Together, these findings provide *in vivo* genetic validation that CAD is a critical determinant of myocardial reperfusion injury. Cardiomyocyte-specific loss of CAD confers robust protection against infarction and promotes functional recovery after I/R.

## Discussion

Myocardial ischemia/reperfusion (I/R) injury arises from the abrupt reintroduction of oxygen and metabolic substrates into a highly reduced cellular environment, triggering a burst of reactive oxygen species (ROS), mitochondrial dysfunction, and cardiomyocyte death.^37,38^ Although metabolic remodeling is recognized as a central determinant of reperfusion injury, the specific biosynthetic pathways that couple metabolic stress to ROS generation and cardiomyocyte injury remain incompletely defined. Here, we identify *de novo* pyrimidine biosynthesis and its rate-limiting enzyme CAD as a previously unrecognized metabolic driver of reperfusion injury. Our findings support a dual-compartment model in which CAD activation simultaneously promotes mitochondrial ROS generation through DHODH-dependent reverse electron transport and cytosolic ROS production through purine insufficiency–linked RNA turnover, thereby amplifying injury through coordinated metabolic and redox mechanisms.

Nucleotide metabolism lies at the intersection of bioenergetics, biosynthesis, and stress adaptation, and its dysregulation contributes to diverse human diseases, including immunodeficiency, neurodevelopmental disorders, cancer, and hyperuricemia.^23,39–41^ However, nucleotide biology has been viewed largely through the lens of energy charge and purine degradation rather than dynamic pathway remodeling in the heart. A major finding of this study is that CAD is rapidly activated during I/R *in vivo* and during sI/R in cardiomyocyte cultures through mTOR–S6K-dependent phosphorylation. Genetic deletion of CAD *in vivo* protects against myocardial I/R injury and improves post-ischemic functional recovery. Consistently, CAD knockdown or pharmacological inhibition *in vitro* mitigates cardiomyocyte injury, whereas CAD activation exacerbates oxidative stress and cell death. These data establish CAD as both necessary and sufficient to modulate susceptibility to reperfusion injury.

Mitochondrial ROS production during early reperfusion has been increasingly recognized as a critical determinant of injury, with complex I reverse electron transport (RET) emerging as a major source of the rapid superoxide burst that drives mitochondrial permeability transition and cell death.^42^ The prevailing model emphasizes succinate accumulation during ischemia and its rapid oxidation through complex II at reperfusion as the key trigger for RET.^7^ Here, we identify an additional metabolic route that converges on the same redox mechanism. Specifically, CAD activation enhances flux through the CAD–DHODH branch of *de novo* pyrimidine biosynthesis, thereby increasing availability of DHOA, the substrate for DHODH, an inner mitochondrial membrane enzyme that transfers electrons to the CoQ pool during oxidation of DHOA to orotate (OA). CAD activation elevated the CoQH_2_/CoQ ratio and increased mitochondrial ROS production during sI/R, whereas CAD silencing or pharmacological inhibition suppressed these effects. Genetic or pharmacological blockade of DHODH abolished CAD-driven mitochondrial ROS and significantly improved cell survival, demonstrating that DHODH is required for this pathway. Mechanistically, exogenous DHOA further enhanced mitochondrial ROS production and exacerbated cell death, supporting a substrate-driven model in which increased flux through DHODH amplifies mitochondrial oxidative stress. In parallel, co-immunoprecipitation studies revealed robust association among CAD, DHODH, and VDAC3, whereas the phosphorylation-defective CAD S1859A mutant exhibited markedly reduced complex formation. Notably, DHOA restored CAD–DHODH–VDAC3 assembly in the S1859A context, indicating that both CAD activation status and substrate availability cooperatively regulate formation of this mitochondrial metabolic signaling complex. Together, these findings establish CAD as an upstream activator of DHODH-dependent RET, broaden prevailing models of reperfusion-associated mitochondrial ROS generation, and demonstrate that pyrimidine biosynthetic flux can directly drive mitochondrial oxidative injury during reperfusion.

At the cytosolic level, CAD activation profoundly rewires nucleotide homeostasis. Targeted metabolite analysis together with isotope tracing demonstrated that CAD activation diverts glutamine-derived nitrogen toward *de novo* pyrimidine synthesis while suppressing *de novo* purine biosynthetic flux. A central advance of this study is the identification of purine insufficiency as a direct determinant of RNA stability during reperfusion stress. CAD overexpression accelerated global RNA decay without affecting nascent RNA synthesis, whereas CAD silencing slowed RNA decay. Importantly, direct inhibition of *de novo* purine synthesis through genetic depletion of GART or pharmacological inhibition with lometrexol was sufficient to accelerate RNA decay even in the absence of CAD overexpression, demonstrating that purine depletion itself drives RNA destabilization rather than representing a secondary consequence of CAD signaling. Conversely, replenishing purine pools with adenine or guanine restored RNA stability, suppressed ROS accumulation, and rescued cell viability, establishing purine availability as a key metabolic checkpoint governing RNA homeostasis under stress. Mechanistically, this pathway requires the cytoplasmic RNA exosome component DIS3L, but not the nuclear catalytic subunit DIS3, indicating compartment-specific control of RNA turnover. DIS3L-dependent RNA degradation generates purine catabolites, including hypoxanthine and xanthine, which are subsequently oxidized by XDH to uric acid with concomitant ROS production.^35^ Silencing DIS3L or XDH, inhibiting XDH enzymatically, or blocking purine nucleoside phosphorylase each markedly reduced cytosolic ROS and improved cell survival during sI/R. Together, these findings establish purine insufficiency as an upstream trigger of RNA decay-coupled oxidative injury rather than a secondary consequence of metabolic stress.

Beyond these specific pathways, our data suggest that CAD activation lowers the threshold for cardiomyocyte injury by engaging two spatially distinct ROS hubs: a mitochondrial pathway mediated by DHODH-dependent RET and a cytosolic pathway driven by purine insufficiency, DIS3L-dependent RNA decay, and purine catabolism. This integrated framework provides a mechanistic explanation for the marked vulnerability of CAD-activated cardiomyocytes to reperfusion stress and for the robust protection conferred by CAD loss, purine restoration, or inhibition of DHODH/XDH signaling.

Collectively, our findings position CAD at the intersection of nucleotide metabolism, RNA stability, and mitochondrial redox biology during reperfusion injury. By demonstrating that CAD coordinates both pyrimidine-driven mitochondrial ROS production and purine depletion–driven cytosolic ROS generation, we identify CAD as a central metabolic regulator of cardiomyocyte fate. Therapeutically, targeting CAD itself or selectively modulating its downstream metabolic outputs may represent a promising strategy to reduce myocardial injury and improve outcomes in ischemic heart disease.

## KEY RESOURCES TABLE

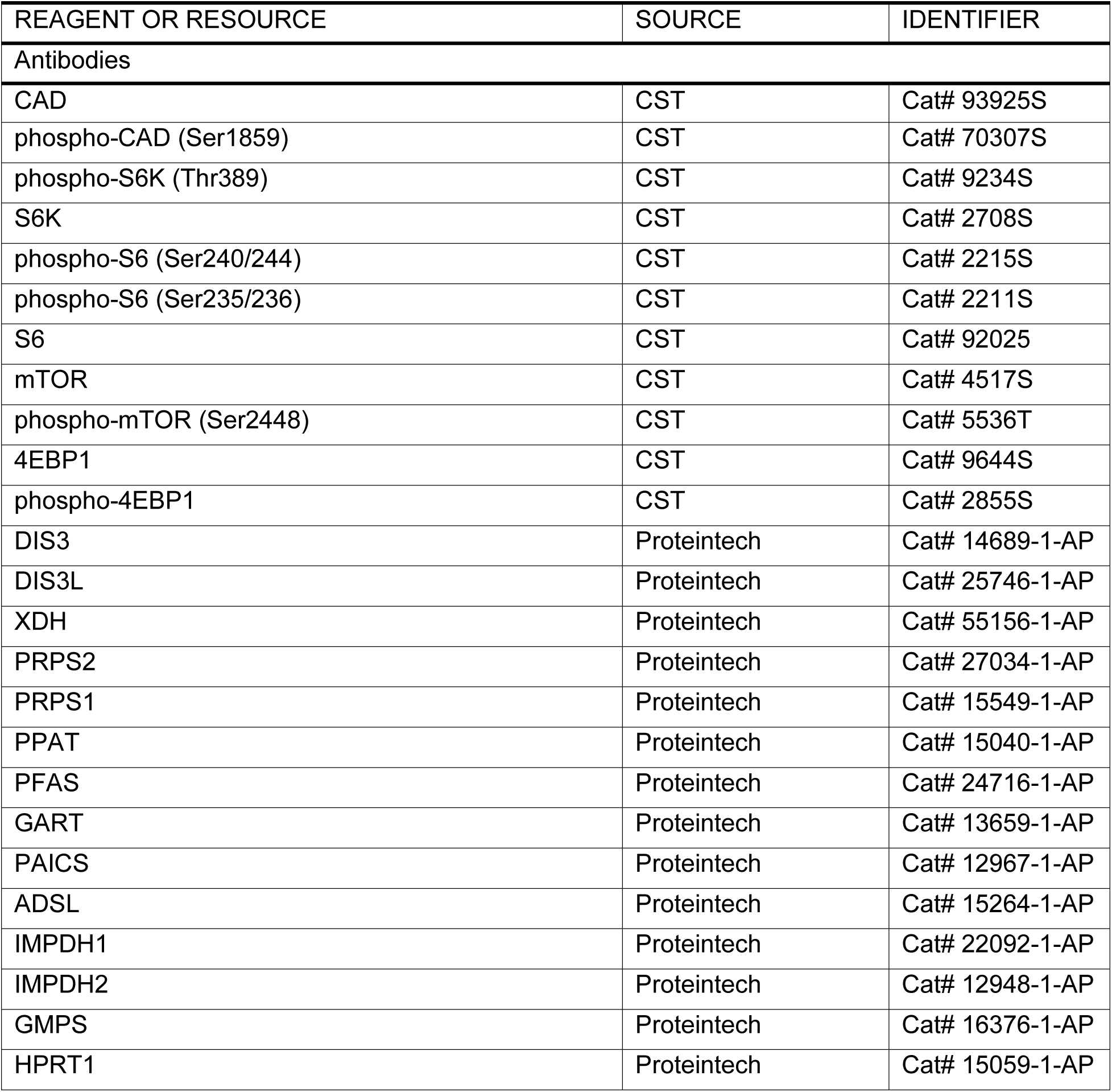
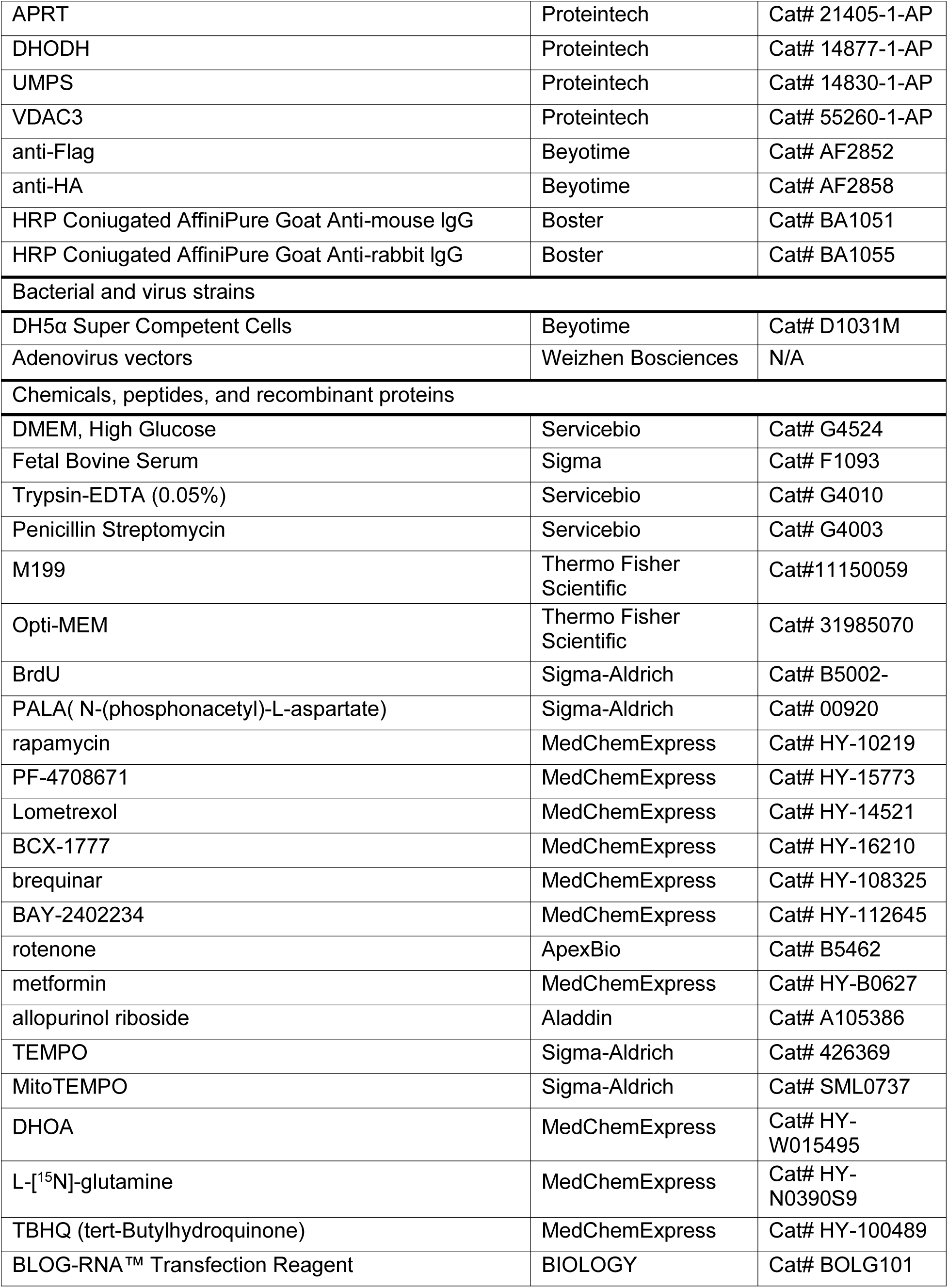
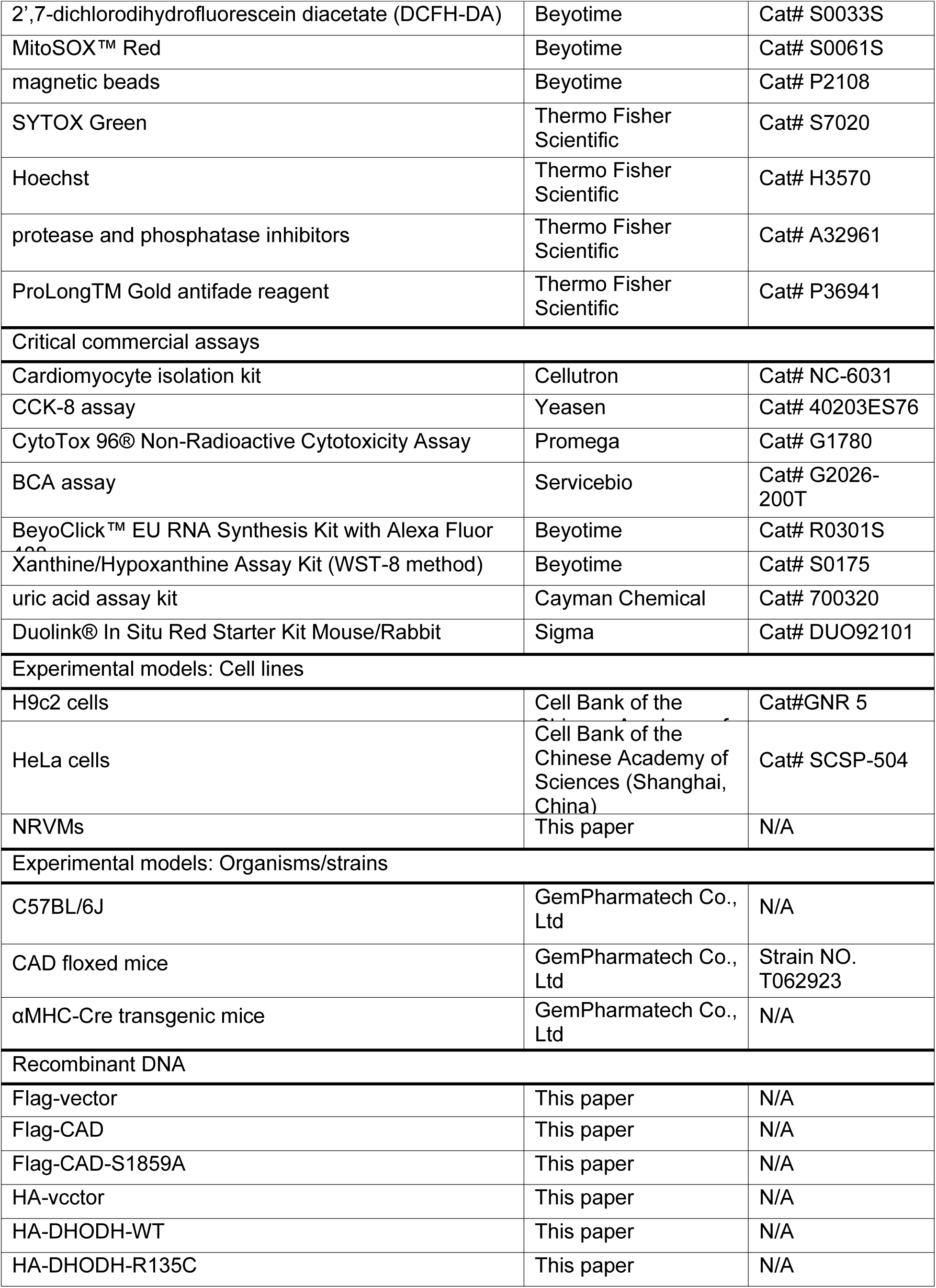
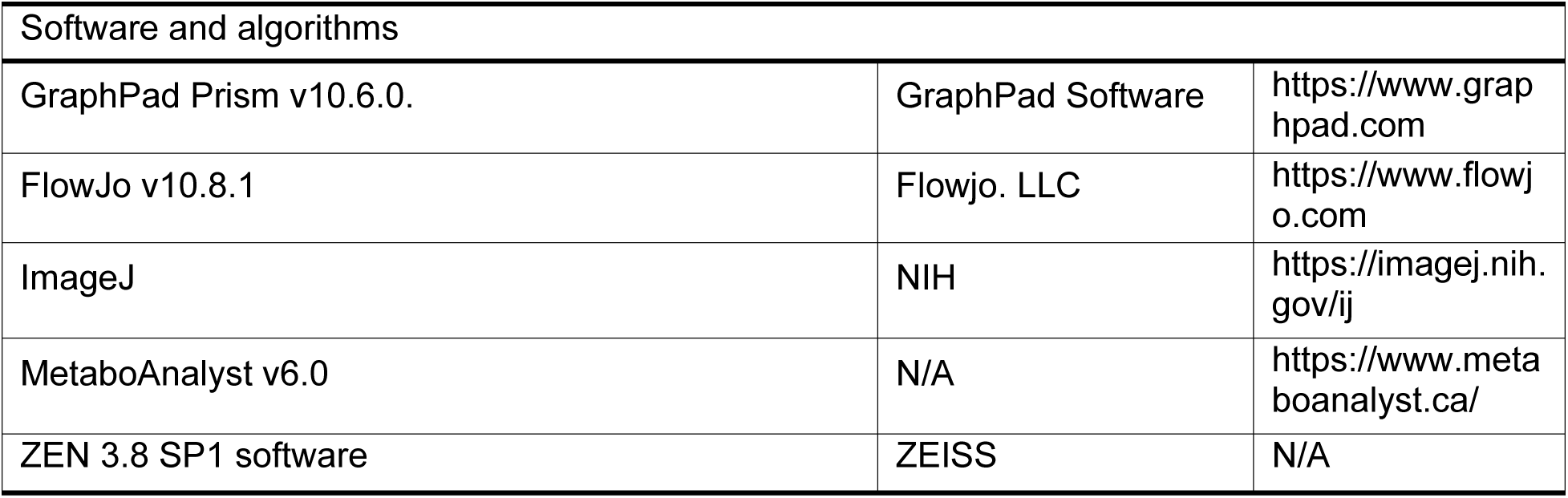

## Methods

### Animals

All animal procedures were approved by the Institutional Animal Research Committee of Tongji Medical College and were performed in accordance with the National Institutes of Health Guide for the Care and Use of Laboratory Animals. Animals were randomly assigned to experimental groups whenever feasible, and investigators performing outcome assessments were blinded to genotype and treatment allocation. Mice were maintained on a C57BL/6 background under controlled conditions with a 12 h light/12 h dark cycle in a temperature-controlled facility and provided ad libitum access to standard chow and water. To generate cardiomyocyte-specific CAD conditional knockout mice, CAD floxed mice were crossed with αMHC-Cre transgenic mice to obtain CAD^fl/fl^; αMHC-Cre mice (CAD cKO). Genotyping was performed by PCR using the following primers: for the CAD floxed allele, forward 5’-GTTCCAATTAAAGGTGCAGCACCAC-3’ and reverse 5’-CCACTTCTACATTATCTACATCATCTCCC-3’; for αMHC-Cre, forward 5’-GATTTCGACCAGGTTCGTTC-3’ and reverse 5’-GCTAACCAGCGTTTTCGTTC-3’.

### Myocardial ischemia/reperfusion (I/R) surgery

Male mice aged 10–14 weeks were subjected to myocardial I/R injury as previously described.^43–45^ Briefly, mice were anesthetized with 4–5% isoflurane for induction and anesthesia was maintained with 1.5%–2% isoflurane in 100% oxygen. Following anesthesia, a small midline cervical incision was made to expose the trachea, and mice were orally intubated for mechanical ventilation. A left thoracotomy was performed via an oblique incision ∼2 mm from the left sternal border, and the fourth intercostal space was opened to expose the heart. After gentle dissection of the pericardium, the left anterior descending (LAD) coronary artery was ligated approximately 1 mm below the tip of the left atrial appendage using a 7-0 polypropylene suture. Successful ischemia was confirmed by pallor of the anterior ventricular wall. After 45 min of ischemia, the ligature was released to allow reperfusion. The chest was closed in layers using absorbable sutures and skin adhesive, and mice were allowed to recover on a warming pad before being returned to their home cages. Sham-operated animals underwent the identical procedure without LAD ligation.

For regional analyses, the left ventricle was subdivided into three regions: the ischemic region, located 1 mm below the ligation site extending toward the apex; the remote region, located 1 mm above the ligature and parallel to the ischemic cut; and the border region, situated between the ischemic and remote zones.

### Echocardiography

Cardiac function was evaluated with unconstrained, conscious mice using high-resolution echocardiography (Visual Sonics, #Vevo 1100, MS400C probe) as previously described.^43,44^ Short-axis M-mode images were obtained at the level of the papillary muscles. Heart rate was recorded. Left ventricular internal diameters at end diastole (LVID, diastolic) and at end systole (LVID, systolic) were measured from M-mode tracings. Fractional shortening (FS) = ((LVID, diastolic - LVID, systolic)/LVID, diastolic) ×100%.

### Chemicals

Where indicated, cells were treated with the following pharmacological agents at the specified final concentrations: PALA (10 µM; CAD inhibitor; N-(phosphonacetyl)-L-aspartate), rapamycin (10 nM; mTORC1 inhibitor; sirolimus), PF-4708671 (5 µM; S6K inhibitor), Lometrexol (5 µM; GART inhibitor), BCX-1777(5 µM; PNP inhibitor), brequinar (10 nM; DHODH inhibitor), BAY-2402234 (10 nM; DHODH inhibitor), rotenone (5 nM; mitochondrial complex I inhibitor), metformin (2 µM; mitochondrial complex I modulator), allopurinol riboside (20 µM; xanthine dehydrogenase inhibitor), TEMPO (10 µM; cytosolic ROS scavenger), and MitoTEMPO (10 µM; mitochondria-targeted superoxide scavenger). DHOA (50 µM).

### Cell culture and siRNA transfection

H9c2 cells and HeLa cells were obtained from the Cell Bank of the Chinese Academy of Sciences (Shanghai, China). H9c2 cells were cultured in high-glucose DMEM supplemented with 10% fetal bovine serum (FBS) and 1% penicillin–streptomycin at 37°C in a humidified incubator with 5% CO_2_. HeLa cells were maintained under identical conditions.

Small interfering RNAs (siRNAs) targeting CAD, DHODH, DIS3L, DIS3, XDH, GART, APRT1, and HPRT1, as well as non-targeting control siRNAs, were purchased from Sigma or Sangon Biotech. siRNAs were resuspended in Opti-MEM (Thermo Fisher Scientific, #31985070) to 40 μM stock concentrations. For transfection, cells were plated in 6-well plates and transfected with ∼80 pmol siRNA per well using BLOG-RNA™ Transfection Reagent (BIOLOGY, BOLG101) according to the manufacturer’s instructions. After 6 hours of incubation, an equal volume of freshly prepared complete medium was added to the culture. All siRNA sequences used in this study are provided in Supplementary Table S1.

### Neonatal rat ventricular myocyte (NRVM) isolation and treatment

NRVMs were isolated from ventricles of 1–2 day-old Sprague-Dawley rats using a commercial cardiomyocyte isolation kit (Cellutron, #NC-6031) as previously described.^44–47^ After 2 hours of pre-plating to remove neonatal fibroblasts, neonatal cardiomyocytes were seeded at a density of ∼1,250 cells mm^-2^ in plating medium consisting of DMEM/M199 (3:1), 5% FBS, 10% horse serum, 1% penicillin-streptomycin, and 100 µM bromodeoxyuridine (BrdU). BrdU was used to suppress neonatal fibroblast growth. After 24 h, medium was replaced with reduced-serum medium (DMEM/M199, 3:1, 1% FBS, 1% penicillin–streptomycin, 100 µM BrdU). After an additional 24 h, cells were switched to serum-free medium prior to experimental treatments, including siRNA knockdown, adenovirus infection, and sI/R.

### Adenovirus production and infection

Recombinant adenoviruses expressing wild-type CAD, phosphorylation-deficient CAD mutants, or empty vector/GFP control were generated by Weizhen Biotechnology (Shandong, China). Cells were infected at the indicated multiplicity of infection (MOI) for 24 h before subsequent experimental treatments.

### Simulated ischemia/reperfusion (sI/R)

Simulated ischemia/reperfusion (sI/R) was performed as previously described.^43–45^ Briefly, H9c2 cells were first washed twice with PBS and incubated in ischemic Esumi buffer (4 mM HEPES, 117 mM NaCl, 0.9 mM CaCl_2_, 0.49 mM MgCl_2_, 12 mM KCl, 5.6 mM 2-deoxyglucose, 20 mM sodium lactate, pH 6.2)^48^. Cells were placed in a modular hypoxia chamber (Billups-Rothenberg, #MIC-101), flushed with 95% N_2_/5% CO_2_ for 30 min (<2 psi), sealed, and incubated at 37 °C for 6-8 h. Reperfusion was initiated by replacing ischemic buffer with complete DMEM and returning cells to normoxic conditions (5% CO_2_). Normoxic control buffer contained 4 mM HEPES, 137 mM NaCl, 0.49 mM CaCl_2_, 0.49 mM MgCl_2_, 0.9 mM KCl, and 5.6 mM glucose (pH 7.4).

Cell viability in H9c2 cells was assessed using the CCK-8 assay (Yeasen, #40203ES76). NRVMs underwent similar sI/R conditions except with a 6-h ischemic period, and cell death was quantified by lactate dehydrogenase (LDH) release (CytoTox96, Promega, #G1780).

### Immunoblotting

Total protein was extracted from cultured cells or cardiac tissue using RIPA lysis buffer supplemented with protease and phosphatase inhibitors (Thermo Fisher Scientific, #A32961). Protein concentration was determined using a BCA assay (Servicebio, #G2026-200T). Equal amounts of protein were loaded on 15-well Criterion SDS-PAGE gels and transferred to nitrocellulose membranes (Bio-Rad, #1704157). Membranes were blocked in 5% BSA for 1 h at room temperature and incubated with primary antibodies overnight at 4°C. Membranes were washed six times (5 min each) in TBST, incubated with HRP-conjugated secondary antibodies for 1 h at room temperature, washed again six times, and visualized using a Tanon 5200 imaging system.

### Immunofluorescence microscopy

Cells were seeded onto glass coverslips in 12-well plates (Biosharp) and allowed to adhere for at least 24 h before experimentation. Following sI/R, cells were fixed with freshly prepared 4% (w/v) paraformaldehyde in PBS for 10 min at room temperature, permeabilized with 0.5% (v/v) Triton X-100 in PBS, and blocked with 1% bovine serum albumin (BSA) in PBS for 1 h at room temperature. Cells were then incubated with primary antibodies overnight at 4°C. After three washes with PBS, samples were incubated with fluorophore-conjugated secondary antibodies for 1 h at room temperature in the dark. Following three additional washes with PBS, slides were mounted with ProLong™ Gold Antifade Reagent (Thermo Fisher Scientific, #P36941). Images were acquired using a ZEISS LSM900 Airyscan confocal microscope and analyzed with ZEISS ZEN 3.8 SP1 software.

### Co-immunoprecipitation

Cells were lysed in ice-cold IP buffer (20 mM Tris-HCl, 100 mM NaCl, 1 mM EDTA-2Na, 10% glycerol, 0.1% NP-40, 1% Triton X-100, pH 7.2) supplemented with protease and phosphatase inhibitors (Thermo Fisher Scientific, #88669). Lysates were clarified by centrifugation at 12,000 × g for 15 min at 4°C. Supernatants were pre-cleared with magnetic beads (Beyotime, P2108) for 1 h at 4°C and then incubated with antibodies and magnetic beads overnight at 4°C with rotation. Beads were washed four times with IP buffer and eluted by boiling in 2× SDS sample buffer for 10 min and analyzed by immunoblotting. For Flag co-IP, anti-Flag (Beyotime, AF2852) were used. For HA co-IP, anti-Flag (Beyotime, AF2858) were used.

### SYTOX Green/Hoechst double staining

SYTOX Green (Thermo Fisher Scientific, #S7020) and Hoechst (Thermo Fisher Scientific, #H3570) were diluted 1:1000 in DMEM containing 3% FBS to generate the working solution. Following sI/R, culture medium was replaced with SYTOX Green/Hoechst working solution, and cells were incubated at 37°C for 30 min in the dark. Cells were then washed three times with PBS and immediately imaged using a fluorescence microscope (Nexcope, #NIB610-FL). SYTOX Green–positive cells were quantified as a measure of membrane-compromised (dead) cells.

### Flow cytometry analysis of reactive oxygen species

For cytosolic general oxidative stress, intracellular reactive oxygen species were measured using 2’,7-dichlorodihydrofluorescein diacetate (DCFH-DA; Beyotime, S0033S). Cells were incubated with DCFDA (10 μM) in serum-free medium at 37°C for 30 min in the dark, washed with PBS containing 1% (w/v) FBS, filtered through a 40-µm nylon mesh, immediately analyzed by flow cytometry. Fluorescence intensity was detected in the FITC channel and quantified as mean fluorescence intensity (MFI).

Mitochondrial superoxide production was assessed using MitoSOX™ Red (Beyotime, S0061S). Cells were incubated with MitoSOX Red (5 μM) at 37 °C for 30 min protected from light, washed with warm PBS containing 1% (w/v) FBS, and analyzed immediately by flow cytometry. Fluorescence was detected in the PE channel. Data were expressed as mean fluorescence intensity.

### EU Pulse–Chase Labeling Assay

H9c2 cells were labeled with 1 mM 5-ethynyl uridine (EU) for 2.5 h at the onset of reperfusion. Cells were fixed immediately (0 h) with 4% paraformaldehyde (PFA) to assess nascent RNA synthesis. For chase experiments, EU-containing medium was replaced with fresh medium lacking EU but supplemented with 1 µM actinomycin D to block transcription. Cells were fixed 1 and 3 h after chase initiation with 4% PFA. EU detection was performed using the BeyoClick™ EU RNA Synthesis Kit with Alexa Fluor 488 (Beyotime, #R0301S) according to the manufacturer’s protocol. Fluorescence images were acquired using a Nikon Eclipse Ti-E inverted microscope equipped with a 20× air objective and an Andor Zyla 4.2 sCMOS camera. Single-cell fluorescence intensity was quantified using NIS-Elements AR v4.30 software. Background fluorescence was corrected by subtracting the mean extracellular signal from each image.

### Analysis of polar metabolites by ion chromatography-mass spectrometry (IC-MS)

Polar metabolites were extracted from heart tissue using 1 mL ice-cold 80/20 (v/v) methanol/water containing 0.1% ammonium hydroxide and 2 mM ^13^C₃-lactate as an internal standard. Extracts were centrifuged at 17,000 × g for 5 min at 4°C, and supernatants were transferred to fresh tubes and dried in a SpeedVac concentrator. Dried extracts were reconstituted in deionized water, and 10 μL was injected for IC–MS analysis.

Ion chromatography was performed on a Thermo Scientific Dionex ICS-5000+ system equipped with an IonPac AS11 column (250 × 2 mm, 4 μm) maintained at 30°C, with the autosampler tray cooled to 4°C. Mobile phase A was water and mobile phase B was 100 mM KOH in water. The flow rate was 360 μL/min with the following gradient: 0–5 min, 1% B; 5–25 min, 1–35% B; 25–39 min, 35–99% B; 39–49 min, 99% B; 49–50 min, 99–1% B (total run time 50 min). Methanol was delivered by an external pump and mixed post-column via a low-dead-volume tee to enhance desolvation and sensitivity. MS data were acquired on a Thermo Orbitrap Fusion Tribrid mass spectrometer in negative ESI mode at 240,000 resolution. Raw files were processed in Thermo TraceFinder, and metabolite abundances were normalized to ^13^C₃-lactate lactate and tissue weight.

### LC–MS analysis of nucleotide metabolites in H9c2 cells

To assess nucleotide metabolites in H9c2 cells, ∼2 × 10^6^ H9c2 cells were seeded per 100-mm dish. Metabolites were extracted using 800 μL ice-cold methanol containing uracil-^13^C_4_, ^15^N_2_ as an internal standard, with stainless-steel beads for homogenization. Samples were centrifuged at 15,000 × g for 15 min at 4°C, and 600 μL supernatant was transferred to a new tube. The extraction was repeated twice with an additional 800 μL ice-cold methanol to ensure complete extraction. Combined extracts were dried under a gentle nitrogen stream, and pellets were retained for DNA quantification.

Dried metabolites were reconstituted in 100 μL methanol and transferred to glass inserts for LC–MS analysis^49^. Measurements were performed on a Waters TQ-XS triple-quadrupole mass spectrometer operated in ESI+ mode. Chromatographic separation was achieved using a Waters ACQUITY UPLC BEH Amide column (2.1 × 100 mm, 1.7 μm) at 40°C. Mobile phase A was water containing 10 mM ammonium formate and 0.1% formic acid; mobile phase B was 85% acetonitrile/15% water containing 10 mM ammonium formate and 0.1% formic acid. The flow rate was 0.4 mL/min, injection volume 10 μL, and the gradient was: 95% B (0.0–1.2 min), 95% to 70% B (1.2–8 min), 70% to 50% B (8–9 min), 50% to 95% B (9–10 min), followed by a 3-min re-equilibration at 95% B. Source parameters were: 150°C source temperature, 500°C desolvation temperature, 1000 L/h desolvation gas flow, 3.0 kV capillary voltage, and argon collision gas at 0.15 mL/min. Data were processed using MassLynx v4.2, and metabolite abundances were normalized to uracil-^13^C_4_, ^15^N_2_ and DNA content.

### Metabolic tracing in H9c2 cells

To assess flux through *de novo* pyrimidine and purine biosynthetic pathways during reperfusion, stable isotope tracing experiments were performed using L-[^15^N]-glutamine. Briefly, approximately 2 × 10^6^ H9c2 cells were seeded in 100-mm culture dishes and subjected to sI/R as described above. At the onset of reperfusion, the culture medium was replaced with reperfusion DMEM containing L-[^15^N]-glutamine at a final concentration of 4 mM. After 6 h of reperfusion, polar metabolites were extracted by ice-cold methanol:acetonitrile:water (2:2:1, v/v/v). Cell lysates were sonicated for 2 min in an ice-water bath and centrifuged at 14,000 × g for 5 min at 4°C. Supernatants were collected, dried by lyophilization, and reconstituted in 50 μL methanol:water (1:1, v/v) immediately prior to LC–MS analysis. Metabolite separation and isotopologue analysis were performed using a Thermo Q Exactive Plus hybrid quadrupole–Orbitrap mass spectrometer coupled to a Thermo Vanquish UHPLC system. Metabolites were resolved on a SeQuant ZIC-HILIC column (100 × 2.1 mm, 3.5 μm; Merck) maintained at 45°C. Mobile phase A consisted of 50 mM ammonium formate in water, and mobile phase B consisted of acetonitrile, delivered at a flow rate of 0.10 mL/min. The gradient was programmed from 90% to 50% B over 10 min, followed by re-equilibration to 90% B. The relative abundance of each metabolite was normalized by total peak intensity. The fractional abundance of each isotopologue was calculated by the peak area of the corresponding isotopologue normalized by the sum of all isotopologue areas.

### CoQ and CoQH2 analysis

Coenzyme Q10 (CoQ10) and its reduced form ubiquinol (CoQ10H2) were extracted from cultured cells using a modified version of the method developed by Nagase et al. and Mao et al. ^22,50^. Briefly, cultured cells were homogenized in 800 μL of ice-cold methanol supplemented with 100 μM tert-butylhydroquinone (TBHQ) using stainless-steel beads. Samples were centrifuged at 15,000 × g for 15 min at 4°C, and 600 μL of the supernatant was transferred to a fresh tube. The extraction was repeated once, and supernatants from both extractions were combined and evaporated to dryness under a gentle stream of nitrogen. Dried extracts were reconstituted in 100 μL methanol, and 10 μL was injected for LC–MS/MS analysis.

LC–MS/MS analysis was performed on an AB Sciex Triple Quad 6500+ mass spectrometer (AB Sciex, Framingham, MA, USA) equipped with an electrospray ionization (ESI) source. Analytes were detected in positive ion mode using multiple-reaction monitoring (MRM) transitions as follows: CoQ10 (m/z 880.7 to 197.1 and 179.1) and CoQ10H2 (m/z 882.7 to 197.1 and 179.1). Chromatographic separation was achieved using a Kinetex C18 column (100 × 2.1 mm, 2.6 μm; Phenomenex) maintained at 40°C. The mobile phase consisted of solvent A (40% water and 60% acetonitrile containing 10 mM ammonium formate and 0.1% formic acid) and solvent B (85% acetonitrile, 10% isopropanol, and 5% water containing 10 mM ammonium formate and 0.1% formic acid), delivered at a flow rate of 0.4 mL min⁻¹. The gradient was as follows: 90% B at 0 min, increased to 100% B at 1.5 min, returned to 90% B at 18 min, and held for 2 min for column re-equilibration. Data acquisition was performed using Analyst software (version 1.7.3), and data processing was conducted using OS software (version 2.1, AB Sciex).

### Xanthine and hypoxanthine measurement

Xanthine and hypoxanthine concentrations in cell were measured using a commercial Xanthine/Hypoxanthine Assay Kit (WST-8 method; Beyotime, #S0175), following the manufacturer’s instructions. For H9c2 cells, cultures were washed once with ice-cold PBS and lysed with 100–200 µL BeyoLysis™ Buffer A for Metabolic Assay per 1 × 10⁶ cells. Lysates were incubated on ice for 5–10 min, centrifuged at 12,000 × g for 3–5 min at 4°C, and supernatants were collected. Absorbance at 450 nm was measured using a microplate reader. Xanthine/hypoxanthine concentrations were calculated based on standard curves and normalized to DNA content to account for differences in cell number.

### Uric acid measurement

Uric acid levels were measured using a commercial uric acid assay kit (Cayman Chemical, #700320) according to the manufacturer’s instructions. Briefly, 105 µL assay buffer, 15 µL uric acid detector (ADHP), and 15 µL sample or standard were added to each well, followed by 15 µL enzyme mixture (uricase and HRP) to initiate the reaction. After incubation at room temperature for 15 min, absorbance at 570 nm was recorded. Background-corrected absorbance values were plotted against a standard curve to determine uric acid concentrations, which were normalized to total DNA content.

### QUANTIFICATION AND STATISTICAL ANALYSIS

All data are presented as mean ± SEM. Comparisons between two groups were performed using two-tailed Student’s *t* test. Comparisons among multiple groups with one variable were analyzed by one-way ANOVA followed by Tukey’s or Dunnett’s multiple-comparisons test. Experiments involving two independent variables were analyzed by two-way ANOVA followed by Tukey’s or Sidak’s post hoc tests. A *P* value < 0.05 was considered statistically significant. Statistical analyses were conducted using GraphPad Prism v10.6.0.

## Acknowledgements

This research was supported by grants from the National Nature Science foundation of China (82470292 to X.W., 82200421 to X.W., 82470407 to G.Z., 82370397 to J.W.), the Noncommunicable Chronic Diseases–National Science and Techonology Major Project of China (2025ZD0547800 to G.Z.), and the National Key Research and Development Program of China (2024YFA1803700 to G.Z. and 2022YFC3400700 to J.W.).

## Author Contributions

Conceptualization, X.Z., S.X. G.Z., and X.W.; Methodology, X.Z., S.X., and X.W.; Investigation, X.Z., S.X., F.Y., Y.S., M.J., Y.Z., K.Z., J.W., F.Z., Y.L., and J.W.; Formal Analysis, X.Z., S.X., G.Z., and X.W.; Resources, J.Wu. G.Z., and X.W.; Writing – Original Draft, X.Z., S.X., and X.W.; Writing – Review & Editing, all authors; Funding Acquisition, J.Wu., G.Z., and X.W.; Supervision, G.Z. and X.W.

